# Common antigenic motif recognized by human VH5-51/VL4-1 tau antibodies with distinct functionalities

**DOI:** 10.1101/287003

**Authors:** Adrian Apetri, Rosa Crespo, Jarek Juraszek, Gabriel Pascual, Roosmarijn Janson, Xueyong Zhu, Heng Zhang, Elissa Keogh, Trevin Holland, Jay Wadia, Hanneke Verveen, Berdien Siregar, Michael Mrosek, Renske Taggenbrock, Jeroen van Ameijde, Hanna Inganäs, Margot van Winsen, Martin H. Koldijk, David Zuijdgeest, Marianne Borgers, Koen Dockx, Esther J.M. Stoop, Wenli Yu, Els C. Brinkman-van der Linden, Kimberley Ummenthum, Kristof van Kolen, Marc Mercken, Stefan Steinbacher, Donata de Marco, Jeroen J. Hoozemans, Ian A. Wilson, Wouter Koudstaal, Jaap Goudsmit

## Abstract

Misfolding and aggregation of tau protein are closely associated with the onset and progression of Alzheimer’s Disease (AD). By interrogating IgG^+^ memory B cells from asymptomatic donors with tau peptides, we have identified two somatically mutated V_H_5-51/V_L_4-1 antibodies. One of these, CBTAU-27.1, binds to the aggregation motif in the R3 repeat domain and blocks the aggregation of tau into paired helical filaments (PHFs) by sequestering monomeric tau. The other, CBTAU-28.1, binds to the N-terminal insert region and inhibits the spreading of tau seeds and mediates the uptake of tau aggregates into microglia by binding PHFs. Crystal structures revealed that the combination of V_H_5-51 and V_L_4-1 recognizes a common Pro-X_n_-Lys motif driven by germline-encoded hotspot interactions while the specificity and thereby functionality of the antibodies are defined by the CDR3 regions. Affinity improvement led to improvement in functionality, identifying their epitopes as new targets for therapy and prevention of AD.

## INTRODUCTION

Intracellular neurofibrillary tangles (NFTs) consisting of aggregated tau protein are a hallmark of Alzheimer’s disease (AD) and other neurogenerative disorders, collectively referred to as tauopathies (Lee et al, 2001). Tau is a microtubule-associated protein expressed predominantly in neuronal axons and promotes the assembly and stability of microtubules (Cleveland et al, 1977b; Weingarten et al, 1975). It is expressed in the adult human brain as six isoforms with zero, one or two N-terminal acidic inserts (0N, 1N, or 2N) and either three or four microtubule-binding repeats (3R or 4R) (Goedert et al, 1989). The tau protein contains many potential phosphorylation sites and the regulated phosphorylation and dephosphorylation of several of these has been shown to affect its interaction with tubulin and cytoskeleton function (Billingsley & Kincaid, 1997; Sanchez et al, 2000). Hyperphosphorylation of tau is thought to lead to microtubule dissociation and assembly of the normally disordered, highly soluble protein into β sheet-rich aggregated fibrils called paired helical filaments (PHFs) that make up NFTs (Cleveland et al, 1977a; Lee et al, 1991; Mandelkow et al, 2007). While the molecular mechanism of tau aggregation remains elusive, it is believed that its initial nucleation step is energetically unfavorable, whereas the subsequent fibril growth follows an energetically downhill landscape (Apetri et al, 2005; Crespo et al, 2012; Harper & Lansbury, 1997; Surewicz et al, 2006). Accumulating evidence indicates that these fibrils can transmit from cell to cell and spread tau pathology to distant brain regions by seeding the recruitment of soluble tau into *de novo* aggregates (Frost et al, 2009; Guo & Lee, 2011; Guo et al, 2016; Kfoury et al, 2012; Wu et al, 2013).

Several monoclonal antibodies that inhibit the spreading of tau fibrils have been described and are being developed for antibody-based therapies (Bright et al, 2015; Umeda et al, 2015; Yanamandra et al, 2013; Yanamandra et al, 2017). We have recently described the isolation of a panel of antibodies with such functional activity by interrogating the peripheral IgG^+^ memory B cells of healthy human blood donors for reactivity to phosphorylated tau peptides (Pascual et al, 2017). To expand the arsenal of potential targets and include epitopes present in physiological tau, we used the BSelex technology in combination with a pool of unphosphorylated tau peptides as baits (Fig. 1A, and table S1 for peptide sequences)

**Fig. 1.**
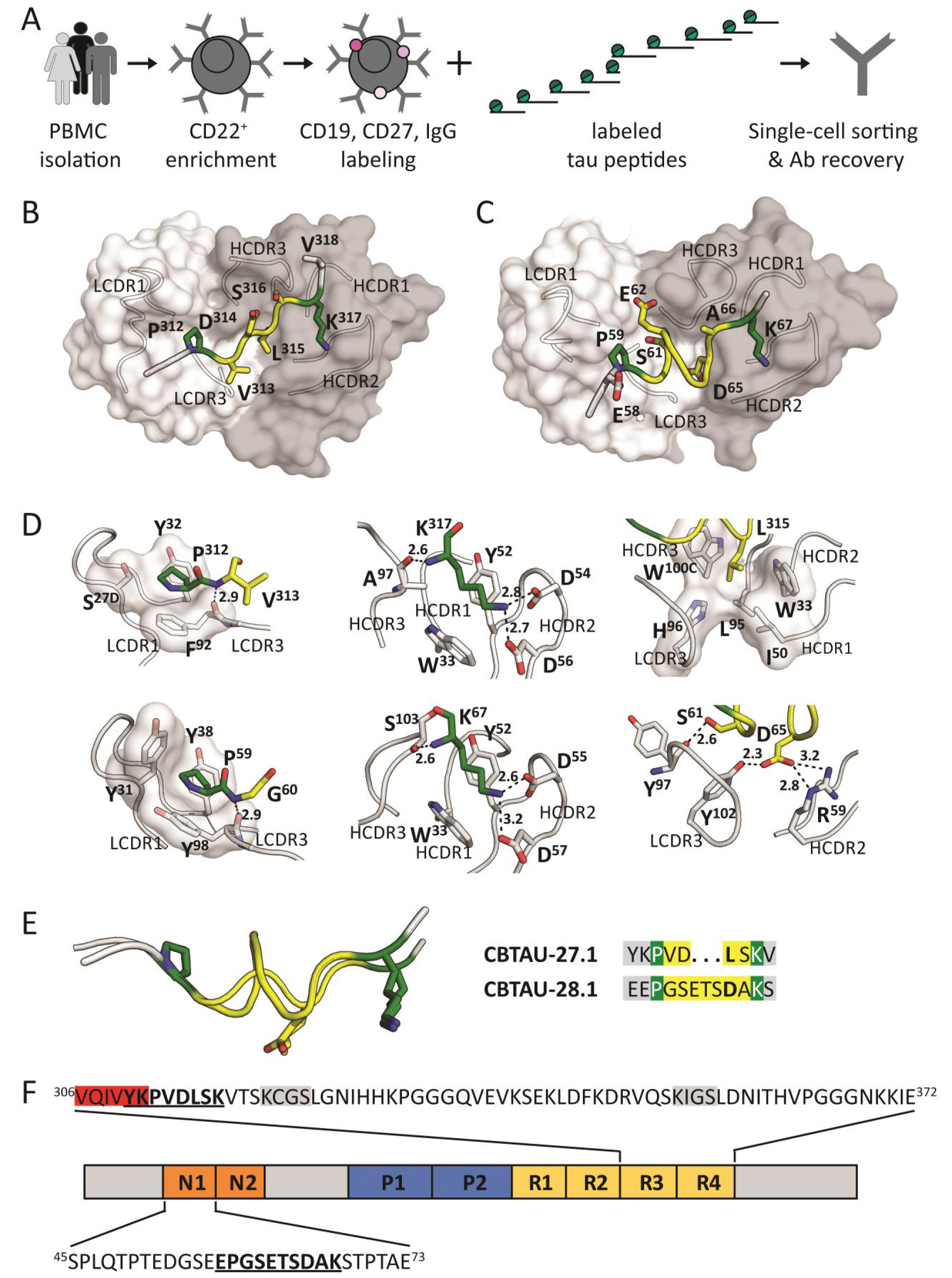
Recovery and structural characterization of naturally occurring monoclonal antibodies to unphosphorylated tau epitopes from asymptomatic individuals. (**A**) BSelex method used to recover tau-specific memory B cells. PBMCs were prepared from asymptomatic blood bank donors, and mature CD22^+^ B cells were positively selected with magnetic beads. Viable cells were stained with IgG-FITC, CD19-PerCPCy5.5, and CD27-PECy7, and with a pool of 10 overlapping unphosphorylated tau peptides spanning the longest tau isoform (relative position of each peptide along 2N4R tau indicated). All peptides were present in the pool with an APC label as well as with a PE label and CD19^+^, CD27^+^, IgG^+^, APC^+^, PE^+^ cells were single-cell sorted on a Beckman Coulter MoFlo XDP. Antibody heavy and light variable chain sequences were recovered from single cells, cloned and expressed as full-length IgGs. (**B and C**) Co-crystal structures of Fab CBTAU-27.1 (**B**) and Fab CBTAU-28.1 (**C**) with tau peptides A8119 and A7731, respectively. Antibodies have been plotted as molecular surface with light chain in white and heavy chain in grey. Tau peptides are shown as cartoon with interacting amino acids plotted as sticks. Proline and lysine residues are plotted in green, amino acids in between these residues are colored in yellow and the termini in grey. Only the interacting antibody loops are outlined. (**D**) Key interactions with tau of CBTAU-27.1 (upper row) and CBTAU-28.1 (lower row). Key interacting residues are plotted as sticks, polar interactions are indicated with dotted lines, and the corresponding distances are indicated in Å. In the first panel, interactions with Pro^312^ and Pro^59^ are compared where the proline binding pockets are visualized on a molecular surface. In the second panel, interactions with Lys^317^ and Lys^67^ are compared. In panel 3, interactions around Leu^315^ and Asp^65^ in the central region of the epitopes are shown. (**E**) Structural basis for recognition of the Pro – X_n_ – Lys epitope motif. Epitopes of CBTAU-27.1 and CBTAU-28.1 are superimposed by aligning on the proline and lysine residues. The peptides present both residues in the same spatial orientation. In the central region (yellow), hydrophobic Leu^315^ in CBTAU-27.1 is replaced by hydrophilic and charged Asp^65^ in CBTAU-28.1. (**F**) Schematic representation of tau isoform 2N4R showing the epitopes of CBTAU-27.1 and −28.1 (bold and underlined) and the surrounding sequences. Highlighted in grey and red are the microtubule binding motifs and the hexapeptide ^306^VQIVYK^311^ which forms the N-terminal end of the core of PHFs, respectively. N1 and N2 indicate acidic inserts, P1 and P2 indicate proline-rich domains, and R1-R4 indicate microtubule-binding repeat domains.

## RESULTS

### Identification of naturally occurring anti-tau antibodies in healthy donors

In total, 2.6 x 10^6^ memory B cells from nine healthy blood donors aged 18-65 years were interrogated and 92 tau-reactive B cells were sorted. From 30 of these, antibody heavy and light variable chain sequences were recovered and full-length IgGs were cloned and expressed. Two unique tau binding antibodies, CBTAU-27.1 and CBTAU-28.1, which are both derived from the V_H_5-51 and V_L_4-1 germline families, were identified. Both antibodies carry high numbers of somatic mutations with 38 and 28 nucleotide substitutions for the heavy and light chains of CBTAU-27.1, and 19 and 16 for the heavy and light chains of CBTAU-28.1, respectively. Since memory B cell selections were performed using a peptide pool, an ELISA-based binding assay with the 10 individual tau peptides was performed and established that CBTAU-27.1 and CBTAU-28.1 bound to peptide A6897 (residues 299-369) and peptide A6940 (residues 42-103), respectively. Further mapping narrowed the epitope regions to residues 299-318 for CBTAU-27.1 and 52-71 for CBTAU-28.1 (fig. S1).

### Recognition of a structurally identical, germline encoded hotspot motif

Crystal structures of the Fab fragments of CBTAU-27.1 and CBTAU-28.1 in complex with tau peptides spanning residues 299-318 and 52-71 were determined at 2.0 and 2.1 Å resolution, respectively (Fig. 1B and C, table S2). The structures reveal that an intriguing similarity exists in the way they bind despite recognition of very distinct regions on the tau protein (Fig. 1D). Both light chains harbor a pocket made of aromatic tyrosine or phenylalanine side chains that form a binding site for a proline residue in the N-terminal region of the different peptides. This interaction is further stabilized by a peptide backbone hydrogen bond to LCDR3 Phe^92^ (CBTAU-27.1) and LCDR3 Tyr^98^ (CBTAU-28.1). Similarly, the two heavy chains interact with a lysine in the peptide C-terminal region that involves identical hydrogen bonding networks with two HCDR2 aspartates and the backbone of HCDR3 Ala^97^ (CBTAU-27.1) and Ser^103^ (CBTAU-28.1), respectively. Both lysines are flanked by HCDR1 Trp^33^ and HCDR2 Tyr^52^ that align and stabilize the aliphatic part of the Lys side chains. However, the central parts of the Tau epitopes differ significantly (see Fig 1D, column 3). The four-residue central region adopts an extended structure in CBTAU-27.1 and inserts Leu^315^ into a pocket formed by LCDR3, HCDR3, HCDR1 and HCDR2. In the same spatial location, the seven-residue central region of the CBTAU-28.1 epitope spans the same distance between the conserved proline and lysine residues by adopting a more compact helical structure (Fig. 1E). CBTAU-28.1 Asp^65^ makes a salt bridge with Arg^59^ (HCDR2) and a hydrogen bond with Tyr^102^ (LCDR3). These two antibodies thus recognize a Pro – X_n_ – Lys motif in different tau peptides, where n is from 4 up to at least 7 amino acids. The proline and lysine binding pockets are germline-encoded and specificity towards one or other epitope arises from the CDR3 loops, which interact with the X region (Fig. 1D).

The CBTAU-27.1 epitope encompasses residues ^310^YKPVDLSK^317^ in the R3 domain (Fig. 1F). The R3 domain is part of the core of PHFs (Daebel et al, 2012; Fitzpatrick et al, 2017), where a hexapeptide ^306^VQIVYK^311^ is crucial for PHF assembly (Daebel et al, 2012; Fitzpatrick et al, 2017; Li & Lee, 2006; von Bergen et al, 2000). Since the CBTAU-27.1 epitope overlaps this hexapeptide, in particular the key Lys^311^ (Li & Lee, 2006), we hypothesize that CBTAU-27.1 binding to tau could block the nucleation interface and thus prevent aggregation. The CBTAU-28.1 epitope encompasses residues ^58^EPGSETSDAK^67^ in the first N-terminal insert (Fig. 1F). Since the N-and C-terminal tau regions that surround the repeat domains have been shown to be disordered and project away from the PHF core to form a flexible fuzzy coat (Fitzpatrick et al, 2017), binding of CBTAU-28.1 is unlikely to interfere with PHF formation, but may—like previously described antibodies (Bright et al, 2015; Umeda et al, 2015; Yanamandra et al, 2013; Yanamandra et al, 2017)—hamper the spreading of aggregates after they are formed. However, the affinities of both antibodies, at least to their cognate tau peptides, are in the high nanomolar range (Fig. S2A and B), which may limit their functional activity. Therefore, we set out to generate affinity-improved mutants of CBTAU-27.1 and CBTAU-28.1 by employing a combination of rational design and random mutagenesis approaches.

### Affinity-improved antibodies retain specificity

For CBTAU-27.1, we used a rational structure-based approach through analysis of the co-crystal structure (Fig. 2A). LCDR3 Thr^94^ was identified as one location where additional hydrophobic interactions could be formed without affecting the structure of the tau peptide. Isoleucine introduced at this position better filled the gap between Val^313^, Leu^315^ and the aliphatic portion of V_H_ Arg^58^. In LCDR1, Ser^27D^ was mutated to tyrosine to remove the unfavorable contact between the serine hydroxyl and the proline pyrrolidine sidechain and create additional hydrophobic interactions. These two mutations improved the affinity by more than 50-fold to the low nanomolar range (Fig. 2C and fig. S2). For CBTAU-28.1, analysis of the structure did not reveal potential affinity-improving mutations and, therefore, a random mutagenesis strategy was employed (Fig. 2B). This approach led to the identification of Ser^32^ →Arg and Glu^35^ → Lys mutations in the light chain that combined led to an ∼4-fold improvement in affinity compared to the parental antibody (Fig. 2D and fig. S2). Co-crystal structures of the Fab fragments of the CBTAU-27.1 double mutant Ser^27D^ → Tyr / Thr^94^ → Ile (from here on referred to as dmCBTAU-27.1) and the CBTAU-28.1 double mutant Ser^32^ → Arg / Glu^35^ → Lys (from here on referred to as dmCBTAU-28.1) in complex with peptides A8119 and A7731, respectively, were determined at 3.0 and 2.85 Å resolution (table S3). Alignment of the structures of the double mutants in complex with their tau epitopes to the corresponding parental antibody co-crystal structures showed that both dmCBTAU-27.1 and dmCBTAU-28.1 retained the binding mode of the parental antibody (Fig. 2E and F), with RMSD values for the peptide Cα atoms of 0.44 Å and 0.24 Å, respectively. The similarity between the double mutants and their parental antibodies regarding the nature of their interactions with tau was confirmed by biolayer interferometry using buffers of different ionic strengths (fig. S3). Furthermore, binding of the different antibodies to sets of tau peptides was assessed to confirm conservation of the specificity (fig. S3).

**Fig. 2.**
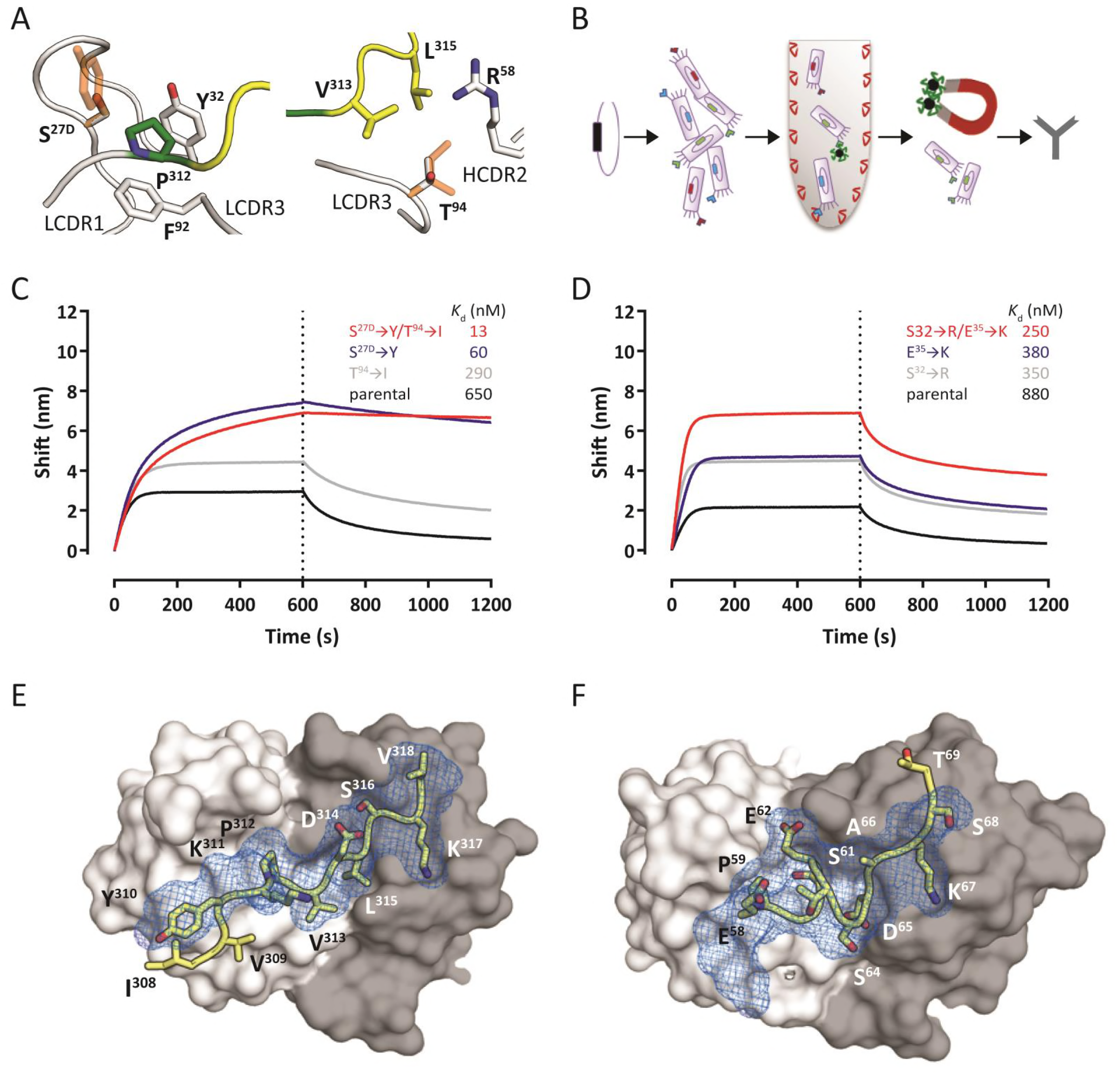
Generation of affinity-improved mutants of CBTAU-27.1 and CBTAU-28.1. (**A**) Structure-based design of mutants around Pro^312^ (left panel) and Val^313^ (right panel). The tau epitope is illustrated as in Figure 1B. Antibody loops and the key residues interacting with Pro^312^ and Thr^94^ are plotted in white. Proposed mutations are shown as orange sticks on top of the corresponding wild-type side chains. Ser^27D^ is mutated to tyrosine (left panel) to enlarge the hydrophobic pocket of Pro^312^, and Thr^94^ is mutated to isoleucine (right panel) to fill the empty cavity surrounding Val^313^ and Leu^315^. By introducing both mutations, additional hydrophobic contacts between Tau and the antibody loops could be formed, potentially resulting in a lower desolvation penalty and increased affinity. (**B**) Schematic representation of the CBTAU-28.1 affinity maturation process by random mutagenesis. Mutations were introduced randomly by error prone PCR in the coding sequence for the single-chain variable fragment (scFv) directed against the CBTAU-28.1 epitope. M13 phage libraries displaying the scFv were screened against rtau and peptide A6940. Affinity-matured variants were identified by phage ELISA and converted into an IgG1 format to assess binding in solution. (**C** and **D**) Association and dissociation profiles for parental and affinity improved CBTAU-27.1 (**C**) and CBTAU-28.1 (**D**) variants to their corresponding cognate peptides as determined by Octet biolayer interferometry. Affinities as determined by ITC (*K*_d_) are shown on the individual graphs. (**E** and **F**) Co-crystal structures of the Fabs of dmCBTAU-27.1 (**E**) and dmCBTAU-28.1 (**F**) with tau peptides A8119 and A7731, respectively. Antibodies are illustrated as molecular surfaces (colored as in panel A), together with tau epitopes as sticks with yellow carbons. The corresponding parental co-crystal structures have been aligned using their variable regions, and their tau epitopes are shown as blue mesh on top of the mutant epitopes.

**Fig. 3.**
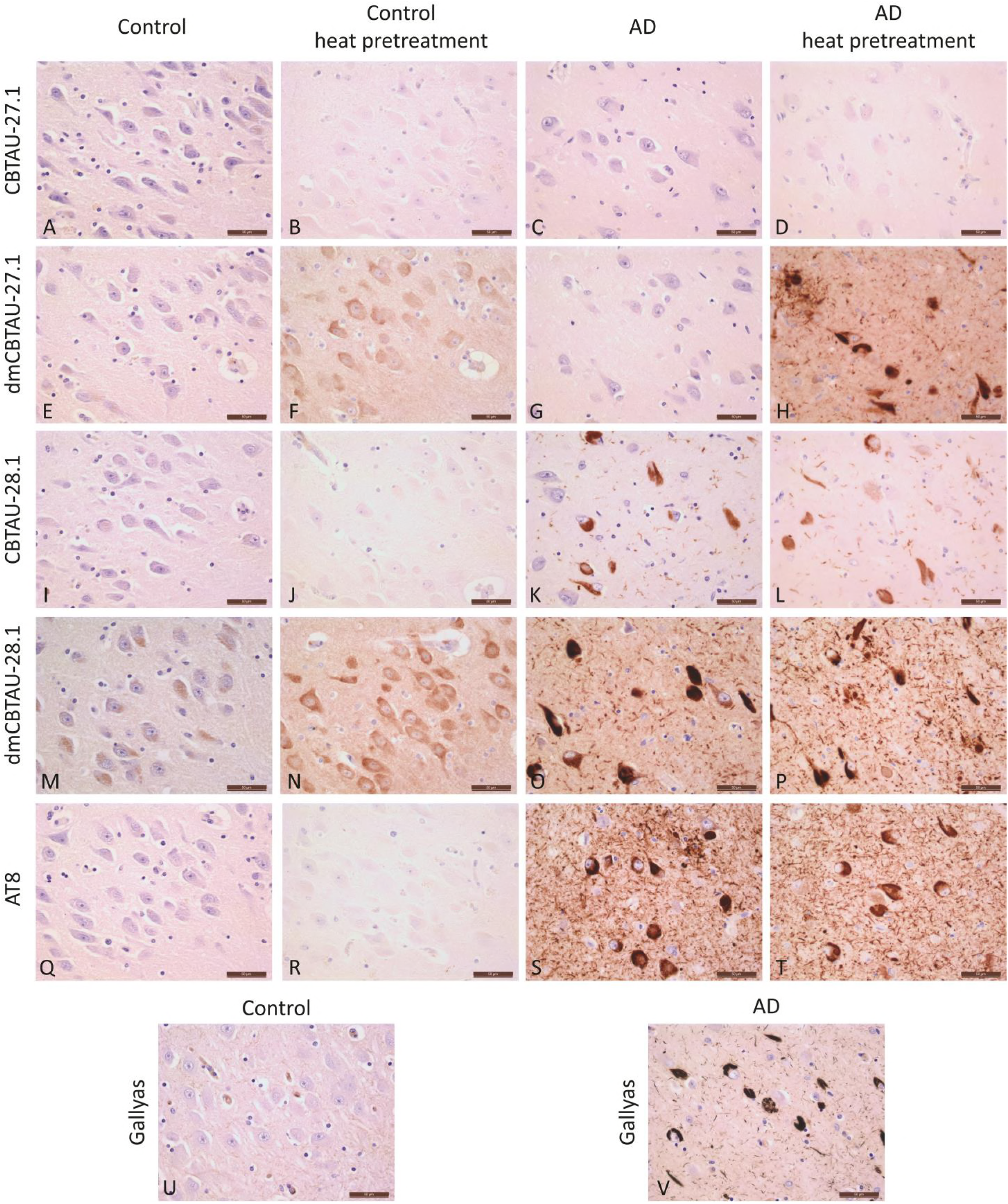
Detection of immunoreactivity in human brain tissue by CBTAU-27.1 and CBTAU-28.1 and affinity-improved variants. Immunohistochemistry was performed on 5 µm thick formalin-fixed paraffin embedded sections of the hippocampal region using a 0.1 µg/ml antibody concentration. Immunodetection using CBTAU-27.1 (**A-D**), dmCBTAU-27.1 (**E-H**), CBTAU-28.1 (**I-L**), dmCBTAU-28.1 (**M-P**) and PHF-tau-specific mouse antibody AT8 (**Q-T**) in control and AD brain tissue without or with heat pretreatment using sodium citrate buffer. Gallyas staining for detection of NFTs and neuropil threads is shown from the same control and AD case of corresponding areas for comparison (**U, V**). Immunoreactivity was visualized using DAB (brown) and nuclei were counterstained with haematoxylin (blue). Representative areas of the CA1/subiculum of the hippocampus are shown. Scale bars represent 50 µm.

The tau specificity of the antibodies was further assessed by immunohistochemical staining on post-mortem control and AD brain tissue (Fig. 3). CBTAU-27.1 did not show immunoreactivity in either control or AD cases, whereas dmCBTAU-27.1 showed immunoreactivity in the cytosol of neurons of the control cases and clear recognition of aggregated tau in neuropil threads and NFTs in AD brains, but only after heat pretreatment Heat pretreatment using sodium citrate buffer is a routine ‘antigen retrieval’ procedure to recover reactivity in formalin-fixed paraffin-embedded tissue sections. No immunoreactivity of CBTAU-28.1 was detected in the control cases, whereas dmCBTAU-28.1 showed diffuse immunoreactivity of neurons after heat pretreatment. In AD brains, both CBTAU-28.1 and dmCBTAU-28.1 recognized neuropil threads and NFTs regardless of the sample treatment. CBTAU-28.1 thus recognizes PHFs without heat pretreatment whereas CBTAU-27.1, even in its high affinity variant, requires heat pretreatment to recognize pathologic tau. This is in line with the epitope of CBTAU-27.1 being buried within the PHFs and becoming exposed upon heat pretreatment. The diffuse neuronal immunoreactivity of dmCBTAU-27.1 and dmCBTAU-28.1 observed in control brain tissue after heat pretreatment shows that these antibodies bind to physiological tau. The observed immunoreactivity under identical conditions shows a clear improvement in the detection of tau by affinity-improved antibodies relative to parental antibodies without affecting specificity (Fig. 3).

### Binding domain-dependent functional activities

The observation that the epitope of CBTAU-27.1 forms part of the core of the PHFs led us to consider that CBTAU-27.1 might be able to prevent aggregation of tau by inhibiting the initial nucleation step. While the molecular mechanism of tau aggregation is not fully understood, the current paradigm is that it follows a nucleation-dependent polymerization (NDP) process (Apetri et al, 2005; Crespo et al, 2012; Harper & Lansbury, 1997; Surewicz et al, 2006). An NDP mechanism is characterized by an initial nucleation step (nuclei formation) followed by an exponential growth step (fibril elongation). Nucleation, the rate-limiting step of the aggregation process, is a stochastic phenomenon and refers to the formation of high energy nuclei. Once nuclei are formed, they rapidly recruit tau monomer (growth step), and convert into thermodynamically stable aggregates. These aggregates can undergo fragmentation generating more fibril ends that are capable of recruiting tau monomers and converting them into *de novo* fibrils. This process is in most general terms referred to as “seeding”. To assess whether CBTAU-27.1 can interfere with the aggregation of tau, we have developed a robust and highly reproducible *in vitro* assay that monitors the heparin-induced aggregation of full-length recombinant Tau441 (rtau) by thioflavin T (ThT) fluorescence. The aggregation behavior of tau in our assay fulfills the expected features of an NDP: sigmoidal kinetic curves with a well-defined lag phase followed by exponential growth ending in a stationary phase (fig. S4). The aggregation kinetics of tau were highly reproducible and displayed the expected concentration dependence (fig. S4B). Furthermore, the obtained tau aggregates displayed PHF-like morphology as assessed by atomic force microscopy (AFM) (fig. S4C) and were extremely efficient in seeding *de novo* aggregation of tau (fig. S4D and E).

**Fig. 4.**
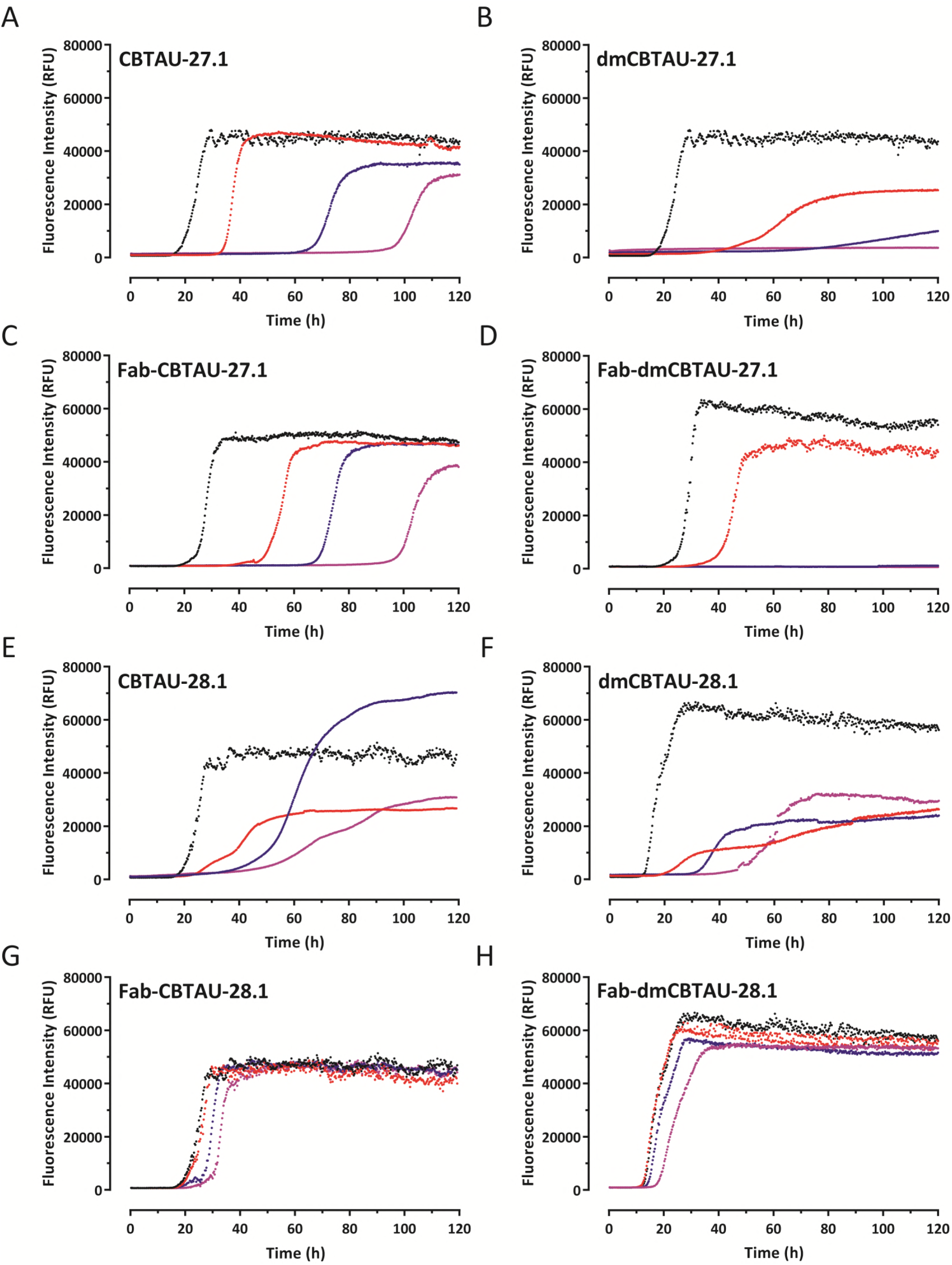
CBTAU-27.1, but not CBTAU-28.1, inhibits the aggregation of recombinant tau *in vitro*. Aggregation of rtau in the absence (black) or presence of CBTAU-27.1 (**A**), dmCBTAU-27.1 (**B**), Fab CBTAU-27.1 (**C**), Fab dmCBTAU-27.1 (**D**), CBTAU-28.1 (**E**), dmCBTAU-28.1 (**F**), Fab CBTAU-28.1 (**G**) or Fab dmCBTAU-28.1 (**H**), as monitored continuously for 120 hr by ThT fluorescence. Three different rtau-IgG (1:0.2 – red, 1:0.4 – blue, and 1: 0.6 – purple) and rtau-Fab (1:0.4 – red, 1: 0.8 – blue, and 1:1.2 – purple) stoichiometries were tested. Each condition was tested in quadruplicate and one representative curve is shown for each condition. For complete datasets, see figs. S5-S8.

In agreement with our hypothesis, CBTAU-27.1 inhibited tau aggregation, as reflected by longer lag phases and lower final ThT fluorescence signal in the kinetic curves. This inhibitory effect was strongly enhanced after affinity improvement (Fig. 4, A and B, and fig. S5). To shed further light on the mechanism, we assessed the ability of dmCBTAU-27.1 to alter the tau conversion when added at later times after initiating the aggregation. In all cases, our results show that stoichiometric amounts of dmCBTAU-27.1 can not only prevent tau aggregation, but also arrest it even in the exponential phase where significant amounts of seeds are already present, presumably by binding and sequestering monomeric tau (fig. S9). The hypothesis that CBTAU-27.1 targets monomeric tau and does not interact with tau aggregates was further confirmed by sedimentation experiments followed by SEC-MALS size determination, which indicates that the antibody binds to two tau monomers using its two Fab arms while not being able to co-sediment with preformed tau aggregates (fig. S10). In sum, these results confirm the initial hypothesis that an antibody that targets the key PHF interface of the monomeric tau can block its misfolding and aggregation.

**Fig. 5.**
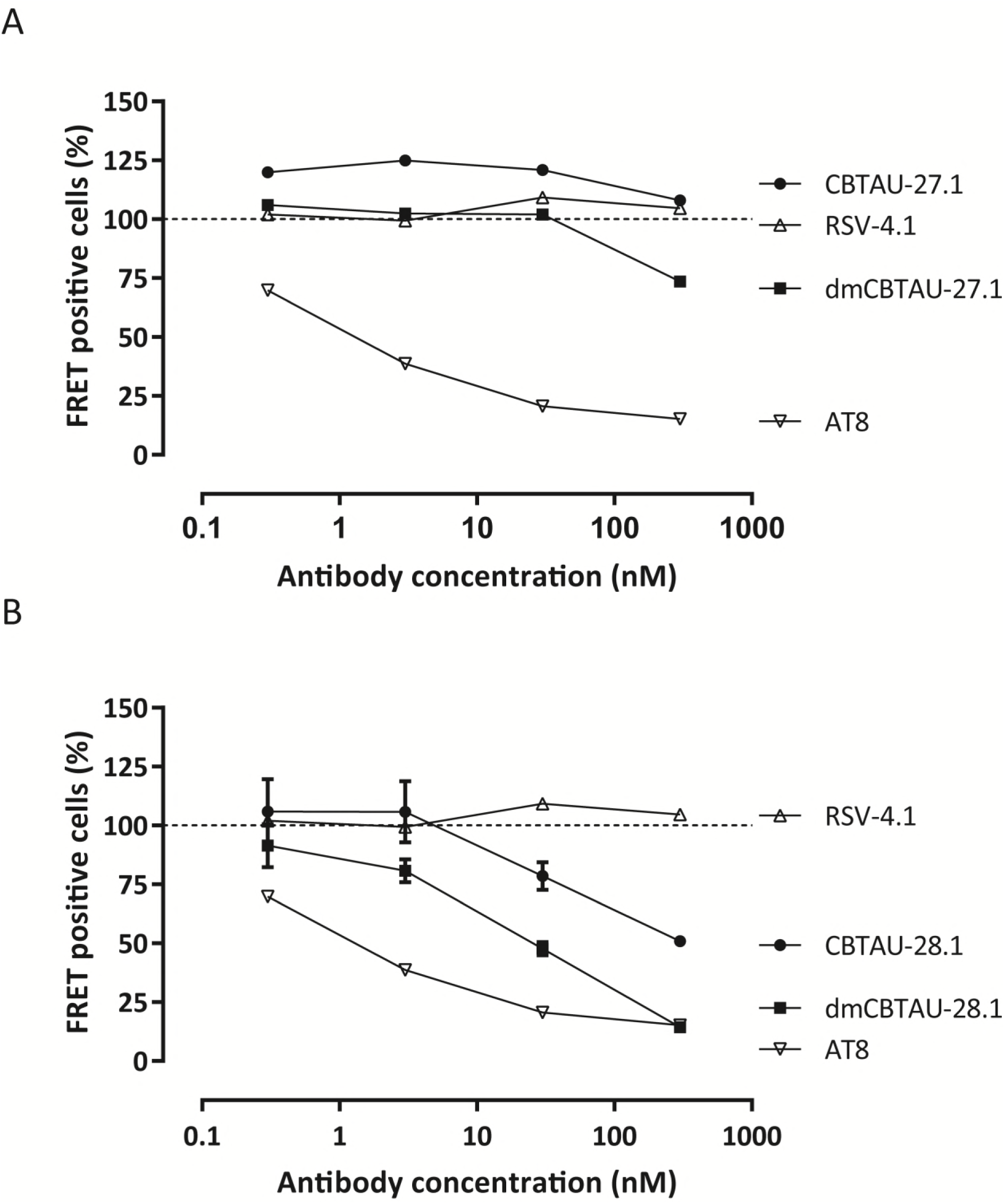
CBTAU-28.1, but not CBTAU-27.1, is capable of immunodepleting seeds from AD brains. Residual seeding activity of human AD brain homogenates following immunodepletion with different concentrations of CBTAU-27.1 and dmCBTAU-27.1 (**A**) or CBTAU-28.1 and dmCBTAU-28.1 (**B**) as measured by FRET signal in biosensor cells expressing the microtubule repeat domains of tau (aa 243-375) fused either to yellow or cyan fluorescent protein. Uptake of exogenous tau aggregates into the cells results in aggregation of the tau fusion proteins, which is detected by FRET. As positive and negative controls, a human IgG1 chimeric version of murine anti-PHF antibody AT8 and anti-RSV-G antibody RSV-4.1 were taken along, respectively. For the controls, the same data are shown in plots A and B for visualization purposes. Error bars indicate the SD of two independent experiments.

Both double mutant and parental CBTAU-28.1 showed comparable alterations in the aggregation kinetics (Fig. 4, E and F, and fig. S6). While somewhat longer lag phases and lower end-point fluorescent signals were observed in the presence of antibody, the effect did not seem to be dose dependent and the kinetics were strikingly irreproducible. Furthermore, visual inspection of reaction mixtures after 120 h incubation revealed that CBTAU-28.1 induced formation of large polymeric structures (fig. S11), suggesting it can crosslink tau aggregates. This notion is supported by the fact that the Fabs of both parental and dmCBTAU-28.1 did not affect tau aggregation (Fig. 4, G and H, and fig. S8). In contrast, CBTAU-27.1 and dmCBTAU-27.1 Fabs showed similar inhibitory effects as their corresponding antibodies (Fig. 4, C and D, and fig. S7), emphasizing the different mechanisms by which CBTAU-27.1 and CBTAU-28.1 interfere with tau aggregation.

**Fig. 6.**
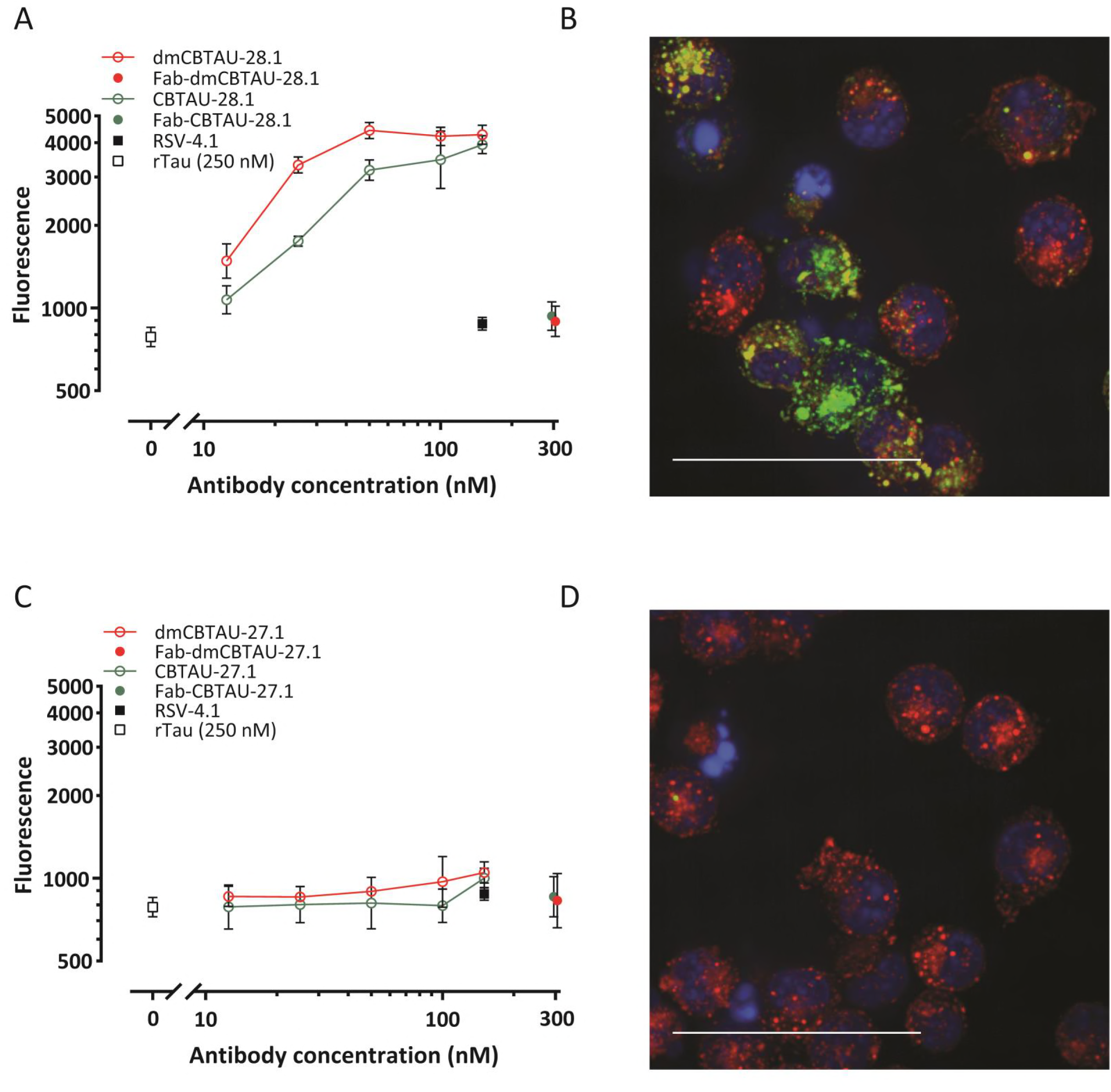
CBTAU-28.1, but not CBTAU-27.1, enhances uptake of tau aggregates into microglial BV2 cells. (**A** and **C**) Aggregated recombinant tau was covalently labelled with pHrodo Green dye and incubated with chimeric versions (containing mouse instead of human Fc region) of CBTAU-28.1, dmCBTAU-28.1, CBTAU27.1, dmCBTAU-27.1, their Fab fragments, a mouse IgG1 isotype control antibody (IC), or no antibody (rtau). Immunocomplexes were subsequently incubated with BV2 cells and their uptake was assessed by flow cytometry as expressed by the geometric mean fluorescent intensity. Error bars indicate the SD of two independent experiments. (**B** and **D**) Preformed pHrodo-Green labeled immunocomplexes of rtau with chimeric dmCBTAU-28.1 or dmCBTAU-27.1 (at a concentration of 150 nM) were incubated with BV-2 cells. After incubation, nuclei were stained with Hoechst (blue) and the acidic cellular compartment with LysoTracker Red dye and uptake of immunocomplexes was assessed by live-cell imaging. Images represent maximum intensity projections of a 20 planes Z-stack (0.5 μm planes) acquired with a 63x water immersion objective.

We next assessed the ability of the antibodies to bind tau aggregates and thus potentially block the propagation and spreading of tau pathology. We therefore incubated human AD brain homogenate containing PHFs with the antibodies and depleted the antibody-antigen complexes. The residual seeding capacity was assessed using a cell-based biosensor assay (Holmes et al, 2014; Yanamandra et al, 2013). In line with its inability to bind PHFs, CBTAU-27.1 did not reduce the seeding activity of the AD brain homogenate, while dmCBTAU-27.1 showed only minor reduction and only at the highest concentration tested (Fig. 5A). In contrast, CBTAU-28.1, like mouse anti-PHF antibody AT8, depletes seeding activity from AD brain homogenate and this *in vitro* activity is enhanced for dmCBTAU-28.1 (Fig. 5B).

The observation that CBTAU-28.1 and dmCBTAU-28 can bind PHFs led us to explore the possibility that these antibodies may furthermore enhance the uptake of tau aggregates by microglia, the resident macrophage cells of central nervous system (Ginhoux et al, 2013). Indeed, CBTAU-28.1 and dmCBTAU-28.1 promoted the uptake of aggregated rtau into mouse microglial BV2 cells and the affinity-improved antibody appeared to mediate tau uptake to a greater extent (Fig. 6A). The fact that Fabs of both parental and dmCBTAU-28.1 did not increase basal tau uptake indicates that the uptake is Fc mediated. As expected, CBTAU-27.1 and dmCBTAU-27.1 did not show activity in this assay (Fig. 6C). Antibody-mediated tau uptake and localization of rtau aggregates in the endolysosomal compartment by CBTAU-28.1 and dmCBTAU-28.1, but not CBTAU-27.1 and dmCBTAU-27.1, was confirmed by confocal microscopy (Fig.6, B and D, and fig. S12).

## DISCUSSION

### Implications for therapy and vaccines

By binding to the region that is critical for the aggregation of tau and which forms the core of PHFs, CBTAU-27.1 prevents aggregation of tau *in vitro*. This functional activity identifies its epitope as a potential target for immunotherapy and could perhaps allow earlier intervention than antibodies that inhibit the spreading of already formed tau seeds (Bright et al, 2015; Umeda et al, 2015; Yanamandra et al, 2013; Yanamandra et al, 2017). Evidence that interfering with tau aggregation through immunotherapy may be possible is provided by murine antibody DC8E8 which also targets monomeric tau, inhibits tau aggregation *in vitro*, and reduces tau pathology in a murine AD model (Kontsekova et al, 2014). An alternative approach could be the development of a small molecule drug targeting monomeric tau that interacts with the CBTAU-27.1 epitope. This epitope does not overlap with the microtubule binding domains (Fig. 1F), suggesting that such a drug may not interfere with the normal function of physiological tau while preventing its aggregation. Furthermore, the epitope of CBTAU-27.1 is specific for tau, and is not present on MAP2, a microtubule stabilizing protein closely related to tau (Dehmelt & Halpain, 2005). Like previously described antibodies, CBTAU-28.1 may be able to inhibit the spread of tau pathology. In addition, it demonstrated an Fc-dependent enhancement of the uptake of tau aggregates by microglial BV2 cells. It is probably able to enhance the uptake of aggregates because its epitope is distant from the core of the PHFs and remains accessible. Both antibodies are derived from the V_H_5-51 and V_L_4-1 germline families and bind their respective epitopes through hotspot interactions that are remarkably alike, pointing towards a conserved structural motif in tau that could not have been predicted from sequence analysis alone. The motif containing proline and lysine separated by 4 to 7 amino acids, is commonly found in Tau, and appears nine times on 2N4R. The lack of somatic mutations around the hotspot proline and lysine suggests these two key tau residues could be responsible for the initial recognition of the V_H_5-51 and V_L_4-1 germline combination. The same V_H_5-51 or V_L_4-1 hotspot interactions can be separately found in other crystallized antibody complexes (Chen, 2017; Gorny et al, 2011; Liao et al, 2013; Ofek et al, 2014), but the combination of the two germlines could be more prevalent against tau, as they seem to get triggered by the Pro – X_n_ – Lys motif abundantly present on the protein. Identification and characterization of these antibodies may thus be exploited to develop antibody or drug regimens for distinct phases of progression of tau pathology and pave the way towards a peptide-based tau vaccine, by taking advantage of the apparent immunogenicity of the identified motif and presenting both hotspot residues in the right spacing and orientation.

## MATERIALS AND METHODS

### Human PBMC isolation

Whole blood from healthy male and female donors was obtained from the San Diego Blood Bank (ages 18-65 years) after informed consent was obtained from the donors. PBMCs were isolated on Ficoll-Paque Plus (GE Healthcare) and cryopreserved at 50 million cells per ml in 90% FBS and 10% DMSO.

### Single cell sorting and recovery of heavy and light variable light antibody genes from tau-specific memory B cells

Tau-peptide specific B cells were sorted and antibody chains were recovered as previously described (Pascual et al, 2017). Briefly, biotinylated peptides were mixed with streptavidin-APC or streptavidin-PE (Thermo Fisher) at a 35:1 molar ratio and free peptide removed using BioSpin 30 column (Biorad). PBMCs corresponding to 3 donors were thawed and B cells were enriched by positive selection with CD22^+^ magnetic beads (Miltenyi Biotec). B cells were subsequently labeled with extracellular markers IgG-FITC, CD19-PerCPCy5.5 and CD27-PECy7 (all from BD Biosciences) and incubated with labeled peptide baits. Doublets and dead cells were excluded and the CD19^+^, IgG^+^, CD27hi, antigen double-positive cells were collected by single cell sorting into PCR plates containing RT-PCR reaction buffer and RNaseOUT (Thermo Fisher). Heavy and light chain (HC/LC) antibody variable regions were recovered using a two-step PCR approach from single sorted memory B cells using a pool of leader specific (Step I) and framework specific primers (Step II PCR). Heavy and light chain PCR fragments (380-400 kb) were linked via an overlap extension PCR and subsequently cloned into a dual-CMV–based human IgG1 mammalian expression vector (generated in house). Finally, all recovered antibodies were expressed and tested by ELISA against the tau peptides used in the sorting experiments to identify tau specific antibodies.

### Peptide synthesis

All peptides were synthesized at Pepscan BV (Lelystad, The Netherlands) or New England Peptide, Inc. using routine Fmoc-based solid phase strategies on automated synthesizers. After acidic cleavage and deprotection, peptides were purified by reversed phase HPLC, lyophilized and stored as powders at −80 °C. Purity and identity were confirmed by LC-MS.

### Recombinant IgG expression and purification

Human IgG1 antibodies were constructed by cloning the heavy (V_H_) and light (V_L_) chain variable regions into a single expression vector containing the wildtype IgG constant regions. Plasmids encoding the sequences corresponding to human anti-tau mAbs were transiently transfected in human embryonic kidney 293-derived Expi293F^TM^ cells (Thermo Fisher) and 7 days post transfection, the expressed antibodies were purified from the culture medium by MabSelect SuRe (GE Healthcare) Protein A affinity chromatography. IgGs were eluted from the column with 100 mM sodium citrate buffer, pH 3.5 which was immediately buffer exchanged into PBS, pH 7.4 using a self-packed Sephadex G-25 column (GE Healthcare). Mouse chimeric versions of CBTAU-27.1 and CBTAU-28.1 were generated cloning the variable light and heavy chain into an expression vector in which the human Fc was replaced with murine Fc. Fab fragments were obtained by papain digestion of antibodies (Thermo Fisher Scientific), followed by removal of the Fc using MabSelect SuRe resin (GE Healthcare). Each antibody was quality controlled by SDS-PAGE and size exclusion chromatography coupled with multi angle light scattering (SEC-MALS) and was further confirmed for reactivity to cognate tau peptide by Octet biolayer interferometry.

### Recombinant tau (rtau) expression and purification

The gene encoding the human tau-441 isoform (2N4R) extended with an N-terminal His-tag and a C-terminal C-tag and containing the C291A and C322A mutations was cloned into a kanamycin resistant bacterial expression vector and transformed into BL21 (DE3) cells. A 3 ml 2-YT Broth (Invitrogen) preculture containing 25 mg/L kanamycin was inoculated from a single bacterial colony for 6 hours after which it was diluted to 300ml for overnight growth in a 1-liter shaker flask and subsequently diluted to 5 liters in a 10-liter wave bag. Protein expression was induced when the culture reached an OD_600nm_ 1.0 by addition of 2 mM IPTG. Three hours after induction, the bacterial pellets were harvested by centrifugation, and lysed with Bugbuster protein extraction reagent (Merck) supplemented with a protease inhibitor cocktail (cOmplete^TM^ ULTRA tablets EDTA free, Roche). Purification was performed by affinity chromatography using self-packed Ni-Sepharose (GE Healthcare) and C-Tag resin (Thermo Fisher Scientific) columns.

### Reactivity of anti-tau human mAbs to tau peptides by ELISA

Reactivity of recovered mAbs were tested against biotinylated tau peptides as previously described (Pascual et al, 2017). Briefly, tau peptides were captured on streptavidin-coated plates (Thermo Fisher Scientific) at 1 µg/ml in TBS and incubated for 2 h. Goat anti-human Fab (2 µg/ml, Jackson ImmunoResearch) to measure total IgG was used, and bovine actin (1 µg/ml TBS, Sigma) and irrelevant peptide were used to confirm specificity of the purified mAbs. ELISA plates were blocked and purified IgGs were diluted to 5 µg/ml in TBS/0.25% BSA and titrated (5-fold serial dilutions) against peptides. Plates were subsequently washed and goat anti-human IgG F(ab’)2 (Jackson ImmunoResearch) was used at 1:2000 dilution as secondary. Following incubation, plates were washed four times in TBS-T and developed with Sure Blue Reserve TMB Microwell Peroxidase Substrate (KPL) for approximately 2 min. The reaction was stopped by the addition of TMB Stop Solution and the absorbance at 450 nm was measured using an ELISA plate reader.

### Qualitative association and dissociation measurements by Octet biolayer interferometry

The relative binding of the antibodies to tau peptides was assessed by biolayer interferometry (Octet Red 384) measurements (ForteBio) (Concepcion et al, 2009). Biotinylated tau peptides were immobilized on Streptavidin (SA) Dip and Read biosensors for kinetics (ForteBio). Real-time binding curves were measured by applying the sensor in a solution containing 100 nM antibody. To induce dissociation, the biosensor containing the antibody-tau peptide complex was immersed in assay buffer without antibody. The immobilization of peptides to sensors, the association and the dissociation steps, were followed in different ionic strength buffers containing 10 % FortéBio kinetics buffer as assay buffer. The relative association and dissociation kinetic curves were compared to qualitatively assess the efficiency of antibody binding to peptides encompassing different tau epitopes.

### Affinity measurements by Isothermal Titration Calorimetry (ITC)

The affinities of antibodies for their corresponding tau peptides were determined in solution on a MicroCal Auto-iTC200 system (Malvern). Peptides at concentrations of ∼35 μM (CBTAU-27.1), ∼10 μM (dmCBTAU-27.1), ∼30 μM (CBTAU-28.1) and ∼33 μM (dmCBTAU-28.1) were titrated in 20 steps of 2 μl per step, except for dmCBTAU-27.1 where 40 steps of 1 μl were employed, in identical buffers containing 200 μM CBTAU-27.1, 130 μM dmCBTAU-27.1, 205 μM CBTAU-28.1 and 205 μM dmCBTAU-28.1, respectively. The thermodynamic parameters and the equilibrium dissociation constants, Kd, were determined upon fitting the ITC data to a model assuming a single set of binding sites corresponding to an antibody:tau peptide = 1:2 binding model.

### Expression, crystallization, data collection, structure determination, and refinement of CBTAU-27.1 Fab and CBTAU-28.1 Fab

To express Fab fragments for crystallization, the gene sequences coding for the hinge, C_H_2, and C_H_3 of human antibodies CBTAU-27.1 (IgG1*κ*) and CBTAU-28.1 (IgG1*κ*) were removed from the corresponding IgG expression vector and a His_6_ tag was added to the C-terminus of each C_H_1. Both Fabs were produced by transient transfection of FreeStyle 293-F cells and purified using a Ni-NTA column followed by a size exclusion chromatography using a Superdex 75 or Superdex 200 column (GE Healthcare). The purified CBTAU-27.1 and CBTAU-28.1 Fabs were concentrated to ∼10 mg/ml and ∼8 mg/ml, respectively, in 20 mM Tris, pH 8.0 and 150 mM NaCl for crystallization.

Crystallization experiments were set up using the sitting drop vapor diffusion method. Initial crystallization conditions for CBTAU-27.1 Fab and its complex with peptide 2833-1 (HVPGGGSVQIVYKPVDLSKV), and CBTAU-28.1 in complex with peptide W1805 (TEDGSEEPGSETSDAKSTPT-amide) were obtained from robotic crystallization trials using the robotic CrystalMation system (Rigaku) at The Scripps Research Institute. For co-crystallization, CBTAU-27.1 and CBTAU-28.1 Fabs were mixed with peptides 2833-1 and W1805, respectively, in a molar ratio of 1:10 before screening. Crystals of *apo*-form CBTAU-27.1 Fab were obtained at 20 °C from a reservoir solution containing 0.1 M Hepes, pH 7.5, 20% PEG8000, while crystals of CBTAU-27.1 Fab with 2833-1 were grown at 20 °C from 0.1 M Hepes, pH 7.0, 0.1 M KCl, 15% PEG5000 MME. Crystals of CBTAU-28.1 with W1805 were obtained at 20 °C from 0.085 M Tris-HCl, pH 8.5, 0.17 M sodium acetate, 25.5% PEG4000. Before data collection, the crystals of CBTAU-27.1 Fab and CBTAU-28.1 Fab were soaked in the reservoir solution supplemented with 25% (v/v) and 15% (v/v) glycerol, respectively, for a few seconds and then flash-frozen in liquid nitrogen. X-ray diffraction data of CBTAU-27.1 Fab *apo*-form were collected to 1.9 Å resolution at beamline 23ID-D at the Advanced Photon Source (APS). X-ray diffraction data of CBTAU-27.1 Fab with 2833-1 and CBTAU-28.1 with W1805 were collected to 2.0 and 2.1 Å resolution, respectively, at beamline 12-2 at the Stanford Synchrotron Radiation Lightsource (SSRL). HKL2000 (HKL Research, Inc.) was used to integrate and scale the diffraction data (Table S2).

The structures were determined by molecular replacement with Phaser (McCoy et al, 2005) using search models of a human antibody 2-23b3 Fab (PDB ID 3QOS) for CBTAU-27.1 Fab and a human antibody 1-69b3 (PDB ID 3QOT) for CBTAU-28.1 Fab. The models were iteratively rebuilt using Coot (Emsley et al, 2010) and refined in Phenix (Afonine et al, 2012). Refinement parameters included rigid body refinement and restrained refinement including TLS refinement. Electron density for both peptides 2833-1 and W1805 were clear and the peptides were built in the later stages of the refinement. Final refinement statistics are summarized in Table S2.

### Crystallization, data collection, and structure determination of dmCBTAU-27.1 Fab and dmCBTAU-28.1 Fab

For crystallization, the dmCBTAU-27.1 and dmCBTAU-28.1 Fab fragments (in 20 mM HEPES buffer, pH 7.5, 7.55 mM NaCl) were incubated with 4 mM of the respective peptide on ice overnight and concentrated to a final concentration of about 50 mg/ml. The Fab:peptide complexes (dmCBTAU-27.1 Fab with tau peptide A8119 and dmCBTAU-28.1 Fab with tau peptide A7731) were subjected to crystallization screening by sitting drop vapor diffusion testing 2,300 conditions, by mixing 0.1µl protein solution and 0.1µl reservoir, and also varying the protein concentration. The dmCBTAU-27.1 Fab with peptide A8119 was crystallized from 0.10 M sodium citrate buffer, pH 5.0, 20.00 % (w/v) PEG 8K at a concentration of 17 mg/ml. The dmCBTAU-28.1 Fab with peptide A7731 was crystallized from 22 % (w/v) PEG5K MME, 0.10 M HEPES buffer, pH 6.75, 0.20 M KCl at a concentration of 50 mg/ml.

For cryo-protection, crystals were briefly immersed in a solution consisting of 75% reservoir and 25% glycerol. X-ray diffraction data were collected at temperature of 100 K at the Swiss Light Source. Data were integrated, scaled and merged using XDS (Kabsch, 2010). The structure was solved with MOLREP (Vagin, 1997) and refined with REFMAC5 (Vagin et al, 2004). Manual model completion was carried out using Coot (Emsley et al, 2010). The quality of the final model was verified PROCHECK (Laskowski, 1993) and the validation tools available through Coot (Emsley et al, 2010).

The dmCBTAU-27.1 diffraction data were indexed in space group C2 and the dmCBTAU-28.1 data in space group P2_1_2_1_2_1_. Data were processed using the programs XDS and XSCALE (see table S3). The phase information necessary to determine and analyze the structure was obtained by molecular replacement using previously solved structure of dmCBTAU-27.1 and dmCBTAU-28.1 Fabs were used search models. Subsequent model building and refinement was performed according to standard protocols with the software packages CCP4 and COOT. For calculation of R-free, 6.2 % of the measured reflections were excluded from refinement (see table S3). TLS refinement (using REFMAC5, CCP4) resulted in lower R-factors and higher quality of the electron density map for dmCBTAU-27.1-A8119 using five TLS groups. Waters were added using the “Find waters” algorithm of COOT into Fo-Fc maps contoured at 3.0 sigma followed by refinement with REFMAC5 and checking all waters with the validation tool of COOT. The occupancy of side chains, which were in negative peaks in the Fo-Fc map (contoured at −3.0 σ), were set to zero and subsequently to 0.5 if a positive peak occurred after the next refinement cycle. The Ramachandran plot of the final model shows 88.7% (dmCBTAU-27.1) and 88.1% (dmCBTAU-28.1) of all residues in the most favored region, 10.8% (dmCBTAU-27.1) and 11.6% (dmCBTAU-28.1) in the additionally allowed region, and 0.3% (dmCBTAU-27.1) and 0.0% (dmCBTAU-28.1) in the generously allowed region. Residue Ala51(L) is found in the disallowed region of the Ramachandran plot (table S3) for both Fabs, as is frequently found in other Fabs (Arevalo et al, 1993). This was either confirmed from the electron density map or could not be modelled in another more favorable conformation. Statistics of the final structure and the refinement process are listed in table S3.

### Affinity maturation by phage display

The coding sequence for scFv directed against CBTAU-28.1 epitope was cloned into an inducible prokaryotic expression vector containing the phage M13 pIII gene. Random mutations were introduced in the scFv by error prone PCR (Genemorph II EZClone Domain Mutagenesis kit) after which the DNA was transformed into TG1bacteria. The transformants were grown to mid-log phase and infected with CT helper phages that have a genome lacking the infectivity domains N1 and N2 of protein pIII, rendering phage particles which are only infective if they display the scFv linked to the full-length pIII (Kramer et al, 2003). Phage libraries were screened using magnetic beads coated with rtau in immunotubes. To deselect nonspecific binders, the tubes were coated with a tau peptide lacking the CBTAU-28.1 epitope. To ensure maturation against the correct epitope, selection was continued using beads coated with the cognate A6940 peptide. Eluted phages were used to infect XL1-blue F’ *E. coli* XL1-blue F’ which were cultured and infected with CT-helper (or VCSM13) phages to rescue phages used for subsequent selection rounds. After three rounds of panning, individual phage clones were isolated and screened in phage ELISA for binding to rtau and cognate CBTAU-28.1 peptide A6940.

### In vitro tau aggregation assay

Stock solutions of 500 μM thioflavin T (ThT) (Sigma-Aldrich, St Louis, MO, USA) and 55 μM heparin (Mw = 17-19 kDa; Sigma-Aldrich, St Louis, MO, USA) were prepared by dissolving the dry powders in reaction buffer (0.5 mM TCEP in PBS, pH 6.7), and filtered through a sterile 0.22 μm pore size PES membrane filter (Corning, NY, USA) or a sterile 0.22 μm pore size PVDF membrane filter (Merck Millipore, Tullagreen, Cork, IRL), respectively. The concentration of the ThT solution was determined by absorption measurements at 411 nm using an extinction coefficient of 22,000 M^-1^ cm^-1^. The huTau441 concentration was determined by absorption measurements at 280 nm using an extinction coefficient of 0.31 ml mg^-1^cm^-1^.

For spontaneous conversions, mixtures of 15 μM huTau441 in 200 μl reaction buffer containing 8 μM heparin and 50 μM ThT were dispensed in 96-well plates (Thermo Scientific, Vantaa, Finland) that were subsequently sealed with plate sealers (R&D Systems, Minneapolis, MN). For seeding experiments, preformed seeds were added to the wells before sealing the plate. To assess the effect of IgG or Fab on the conversion, huTau441 and IgG or Fab were mixed and incubated for 20 minutes in reaction buffer before the addition of heparin and ThT.

Kinetic measurements were monitored at 37 °C in a Biotek Synergy Neo2 Multi-Mode Microplate Reader (Biotek, VT, USA) by measuring ThT fluorescence at 485 nm (20 nm bandwidth) upon excitation at 440 nm (20 nm bandwidth) upon continuous shaking (425 cpm, 3mm).

### Atomic Force Microscopy

For each sample, 20 μl of rtau solution was deposited onto freshly cleaved mica surface. After 3 minutes incubation, the surface was washed with double-distilled water and dried with air. Samples were imaged using the Scanasyst-air protocol using a MultiMode 8-HR and Scanasyst-air silicon cantilevers (Bruker Corporation, Santa Barbara, USA). Height images of 1024×1024 pixels in size and surface areas of 10×10 µm were acquired under ambient environmental conditions with peak force frequency of 2 KHz.

### Assessment of CBTAU-27.1 binding to rtau and PHFs by SEC-MALS

15 μM monomer or aggregated rtau were incubated with dmCBTAU-27.1 in a rtau:IgG1 = 1:0.6 ratio for 15 minutes and samples were subsequently centrifuged for 15 minutes at 20,000 g. Same procedure was also applied for controls containing only monomer rtau, aggregated rtau or IgG. All samples were analyzed by SEC-MALS upon fractionation on a **TSKgel G3000SWxl (**Tosoh Bioscience) gel filtration column equilibrated with 150 mM sodium phosphate, 50 mM sodium chloride at pH 7.0. at a flow rate of 1 mL/min. For molar mass determination, in-line UV (Agilent 1260 Infinity MWD, Agilent Technologies), refractive index (Optilab T-rEX, Wyatt Technology) and 8-angle static light scattering (Dawn HELEOS, Wyatt Technology) detectors were used.

### Immunohistochemistry

Brain samples were obtained from The Netherlands Brain Bank (NBB), Netherlands Institute for Neuroscience, Amsterdam. All donors had given written informed consent for brain autopsy and the use of material and clinical information for research purposes. Neuropathological diagnosis was assessed using histochemical stains (haematoxylin and eosin, Bodian and/or Gallyas silver stains (Uchihara, 2007) and immunohistochemistry for Amyloid beta, p-tau (AT8), α-synuclein, TDP-43 and P62, on formalin-fixed and paraffin-embedded tissue from different parts of the brain, including the frontal cortex (F2), temporal pole cortex, parietal cortex (superior and inferior lobule), occipital pole cortex, amygdala and the hippocampus, essentially CA1 and entorhinal area of the parahippocampal gyrus. Staging of pathology was assessed according to Braak and Braak for tau pathology and Thal for Amyloid beta pathology (Braak & Braak, 1991; Thal et al, 2002).

Formalin-fixed and paraffin embedded tissue sections (5 μm thick) were mounted on Superfrost Plus tissue slides (Menzel-Gläser, Germany) and dried overnight at 37°C. Sections were deparaffinised and subsequently immersed in 0.3% H_2_O_2_ in phosphate-buffered saline (PBS) for 30 min to quench endogenous peroxidase activity. Sections were either treated in sodium citrate buffer (10 mM sodium citrate, pH 6.0) heated by autoclave (20 min at 130 °C) for antigen retrieval or processed without heat pretreatment. Between the subsequent incubation steps, sections were washed extensively with PBS. Primary antibodies were diluted in antibody diluent (Immunologic) and incubated overnight at 4°C. Secondary EnVison^TM^ HRP goat anti-rabbit/mouse antibody (EV-GαM^HRP^, DAKO) was incubated for 30 min at room temperature (RT). 3,3-Diaminobenzidine (DAB; DAKO) was used as chromogen. Sections were counterstained with haematoxylin to visualize the nuclei of the cells, dehydrated and mounted using Quick-D mounting medium (BDH Laboratories Supplies, Poole, England).

For the Gallyas silver staining, 30 µm thick sections were rinsed in distilled water and incubated in 5% periodic acid for 30 minutes at RT, followed by an incubation in silver iodide solution (4% sodium hydroxide, 10% potassium iodide and 0.35% silver nitrate in distilled water) for 30 minutes at RT. Subsequently, sections were washed in 0.5% acetic acid and developed with developer working solution (10 volumes 5% sodium carbonate solution, 3 volumes solution 0.2% ammonium nitrate, 0.2% silver nitrate and 1% Tungstosilicic acid solution, and 7 volumes 0.2% ammonium nitrate, 0.2% silver nitrate, 1% Tungstosilicic acid and 0.3% formaldehyde solution. After color development, sections were rinsed in 0.5% acetic acid, after which sections were incubated in 5% sodium thiosulphate and rinsed in distilled water. Stained sections were mounted on coated glass slides (Menzel-Gläser) and dried for at least 2 hours at 37°C. Subsequently sections were fixed in ethanol 70% for 10 minutes, counterstained with hematoxylin, dehydrated and mounted with Quick D mounting medium.

### FRET based cellular immunodepletion assay

Cryopreserved brain tissue was acquired from the Newcastle Brain Tissue Resource biobank. Frozen brain tissue samples from 17 AD patients were homogenized in homogenization buffer (10 mM Tris (Gibco), 150 mM NaCl (Gibco) containing protease inhibitors (cOmplete^TM^ ULTRA tablets EDTA free, Roche) to obtain a 10% w/v pooled brain homogenate. Individual antibody dilutions were prepared in PBS pH 7.4 (Sigma), mixed with brain extract in a 1:1 ratio in a 96 well PCR plate (Thermo Scientific), and incubated until the beads were washed. Protein-G DynaBeads (Life Technologies) were added in a 96-well PCR plate (Thermo Scientific) and washed twice with PBS, 0.01% Tween20 (Sigma) by pulling down the beads with a magnet (Life Technologies). Wash buffer was removed completely and 10 µl of PBS, 0.1% Tween20 were added to the beads together with 90 µl of the 1:1 antibody-brain extract mixture. Samples were incubated over night at 4°C, rotating at 5 rpm. The following day, the immunodepleted fractions were separated from the beads by pulling down the beads with the magnet, transferred to a new 96-well PCR plate and stored at −80°C until tested. Each condition was tested in duplicate. Immunodepleted fractions were incubated for 10 minutes with Lipofectamine 2000 (Invitrogen) in Opti-MEM (Gibco) in a 96-well cell culture plate (Greiner Bio-one) before 5.5×10^3^ HEK biosensor cells (provided by M. Diamond, Washington University School of Medicine) were added to each well. After a 2-day incubation at 37°C, cells were washed twice with PBS, detached using Trypsin/EDTA (Gibco) and transferred to a polypropylene round bottom plate (Costar) containing FACS buffer (Hank’s Balanced Salt Solution (Sigma), 1mM EDTA (Invitrogen), 1% FBS (Biowest)). Cells were then analyzed for FRET positivity by flow cytometry using a FACS Canto II (BD Bioscience). Each plate contained a brain extract only condition (to assess baseline FRET response) and an antibody isotype control. Results are reported as normalized values, relative to condition without antibody.

### Microglia assay

Aggregated recombinant 2N4R tau (rtau) was generated in the absence of ThT under the conditions described for the in vitro Tau Aggregation Assay and covalently labelled with pHrodo® Green STP Ester (Invitrogen) following manufacturer’s instructions. Briefly, rtau aggregates were spun down by centrifugation at 20800 rcf for 30 minutes and then resuspended in 0.1 M sodium bicarbonate buffer, pH 8.5, at a final concentration of 2 mg/ml. Efficiency of aggregation was assessed by detecting presence of tau in the supernatant using SEC-MALS. Prior to labeling, Tau aggregates were briefly sonicated. Ten moles of dye were added per mole of protein and the mixture was incubated for 45 minutes at room temperature, protected from light. Unconjugated dye was removed using a PD10 column (GE Healthcare) equilibrated with 0.1 M sodium bicarbonate buffer pH 8.5 and eluting the protein with the same buffer. Eluted fractions were evaluated for their protein content by BCA assay (Thermo Fisher Scientific) following the manufacturer’s instructions. Protein containing fractions were pooled, aliquoted and stored at −20°C.

BV-2 cells were cultured in DMEM supplemented with 10% FBS, 100 U/ml penicillin, 100 µg/ml streptomycin and 2 mM L-Glutamine. Cultures were maintained in humidified atmosphere with 5% CO2 at 37°C.

In order to generate immunocomplexes, 250 nM aggregated rtau, covalently labelled with pHrodo Green dye, was incubated with a serial dilution (12.5 – 150 nM) of a chimeric version (mouse Fc region) of CBTAU-28.1 (parental and high affinity mutant) or CBTAU-27.1 (parental and high affinity mutant) in serum-free medium. Tau immunocomplexes were also generated with 300 nM Fab fragments of both CBTAU-28.1 and CBTAU-27.1, in the parental and high affinity mutant format. In each experiment, a mouse IgG1 isotype control was included together with cells incubated with only aggregated rtau. Immunocomplexes were incubated over night at 4°C and the day after applied to BV2 cells for 2 hours at 37°C with 5% CO_2_. During the incubation, antibody-independent Tau uptake was prevented by blocking the Heparan Sulfate Proteoglycan Receptor with 200µg/ml Heparin. After incubation, cells were harvested with 0.25% trypsin-EDTA for 20 min thus simultaneously removing Tau bound to the extracellular membrane, centrifuged at 400 rcf to remove medium, washed twice with PBS, and resuspended in flow cytometry buffer (PBS 1x plus 0.5% BSA and 2mM EDTA). Cells were analyzed with a Canto II flow cytometer (BD) gating for live single cell population, as identified by forward and side scatter profiles. Results are reported as geometric mean fluorescent intensities. Each experiment was performed twice.

For the microscopy experiments, cells were seeded in 96-well µClear® plate (Greiner Bio-one). After incubation with the immunocomplexes, nuclei were stained with Hoechst (Sigma) and the acidic cellular compartment with LysoTracker Red dye (Thermo Fisher). Live-cell imaging was performed using the Opera Phenix™ High Content Screening System (PerkinElmer) with temperature set to 37°C and in presence of 5% CO_2_. For high quality images, a 63x water immersion objective was used and 0.5 μm planes (20 per Z-stack) were acquired per imaged field.

## ACKNOWLEDGEMENTS

We would like to thank Mohammed Drissi Saidi, Hector Quirante, Başak Kükrer, Otto Diefenbach, Tariq Nahar and Imke Sprengers, for protein generation and analysis, Alberto Carpinteiro Soares and Tjado Morrema for technical assistance, Frederique Bard and Louis de Muynck for valuable comments and advice. Human brain tissue for the immunodepletion experiments performed in this study was provided by the Newcastle Brain Tissue Resource which is funded in part by a grant from the UK Medical Research Council (G0400074), by NIHR Newcastle Biomedical Research Centre and Unit awarded to the Newcastle upon Tyne NHS Foundation Trust and Newcastle University, and as part of the Brains for Dementia Research Programme jointly funded by Alzheimer’s Research UK and Alzheimer’s Society.

---

**Table S1.**
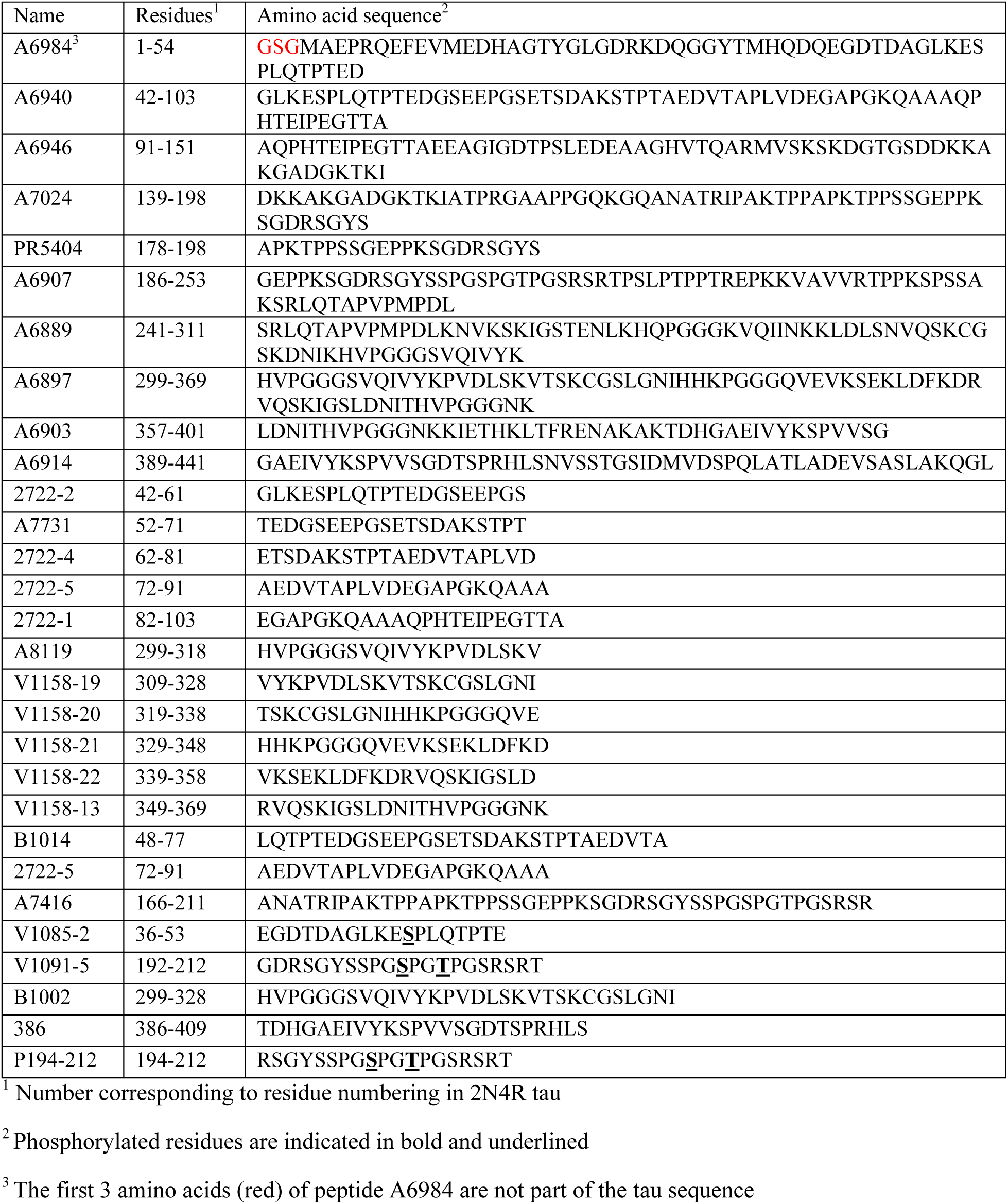
Names and sequences of tau peptides used in this study. The first 10 peptides listed were used as baits in the BSelex method.

**Table S2.**
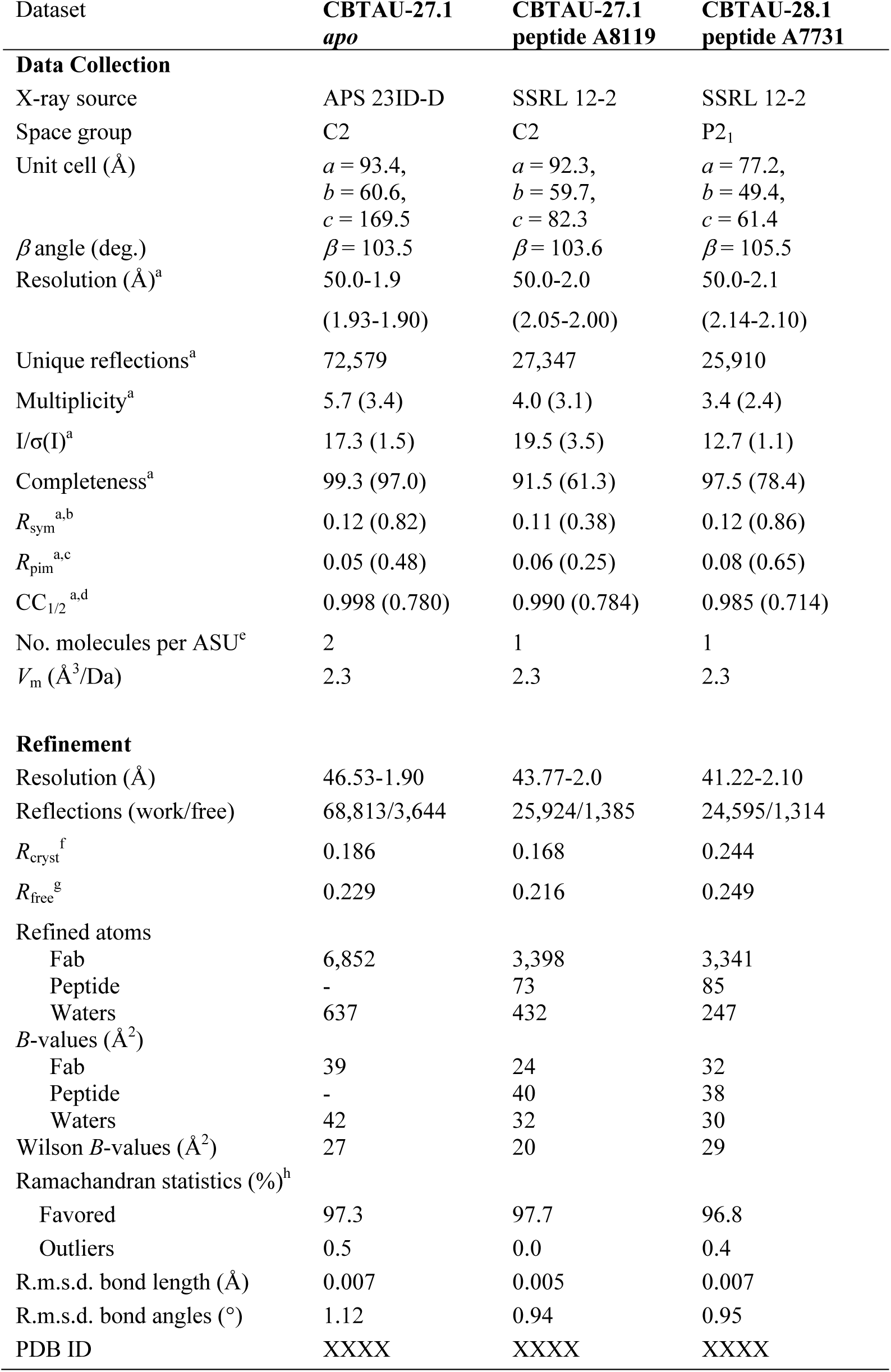
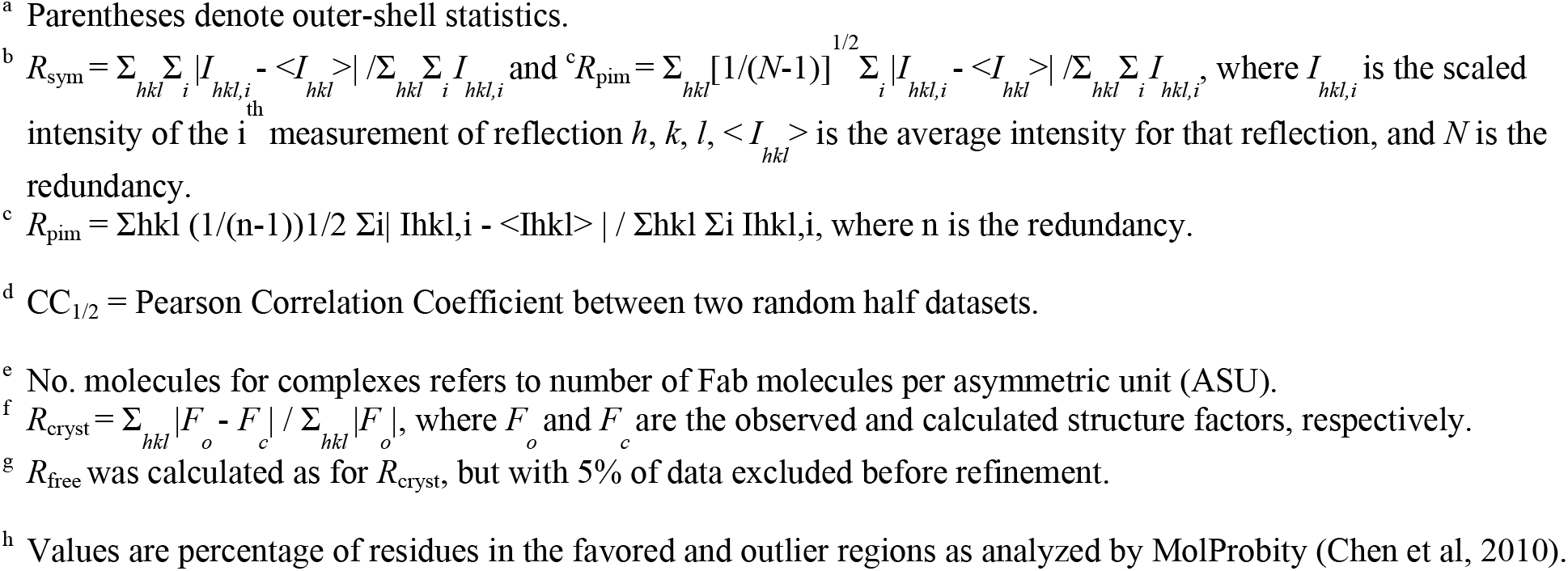
Data collection and refinement statistics for CBTAU-27.1 Fab and CBTAU-28.1 Fab

| Dataset | CBTAU-27.1<br>*apo* | CBTAU-27.1<br>peptide A8119 | CBTAU-28.1<br>peptide A7731 |
| --- | --- | --- | --- |
| **Data Collection** |  |  |  |
| X-ray source | APS 23ID-D | SSRL 12-2 | SSRL 12-2 |
| Space group | C2 | C2 | P2<sub>1</sub> |
| Unit cell (Å) | $a = 93.4,$<br>$b = 60.6,$<br>$c = 169.5$ | $a = 92.3,$<br>$b = 59.7,$<br>$c = 82.3$ | $a = 77.2,$<br>$b = 49.4,$<br>$c = 61.4$ |
| $\beta$ angle (deg.) | $\beta = 103.5$ | $\beta = 103.6$ | $\beta = 105.5$ |
| Resolution (Å)<sup>a</sup> | 50.0-1.9 | 50.0-2.0 | 50.0-2.1 |
|  | (1.93-1.90) | (2.05-2.00) | (2.14-2.10) |
| Unique reflections<sup>a</sup> | 72,579 | 27,347 | 25,910 |
| Multiplicity<sup>a</sup> | 5.7 (3.4) | 4.0 (3.1) | 3.4 (2.4) |
| I/$\sigma$(I)<sup>a</sup> | 17.3 (1.5) | 19.5 (3.5) | 12.7 (1.1) |
| Completeness<sup>a</sup> | 99.3 (97.0) | 91.5 (61.3) | 97.5 (78.4) |
| $R_{\text{sym}}$<sup>a,b</sup> | 0.12 (0.82) | 0.11 (0.38) | 0.12 (0.86) |
| $R_{\text{pim}}$<sup>a,c</sup> | 0.05 (0.48) | 0.06 (0.25) | 0.08 (0.65) |
| $CC_{1/2}$<sup>a,d</sup> | 0.998 (0.780) | 0.990 (0.784) | 0.985 (0.714) |
| No. molecules per ASU<sup>e</sup> | 2 | 1 | 1 |
| $V_m$ (Å<sup>3</sup>/Da) | 2.3 | 2.3 | 2.3 |
| **Refinement** |  |  |  |
| Resolution (Å) | 46.53-1.90 | 43.77-2.0 | 41.22-2.10 |
| Reflections (work/free) | 68,813/3,644 | 25,924/1,385 | 24,595/1,314 |
| $R_{\text{cryst}}$<sup>f</sup> | 0.186 | 0.168 | 0.244 |
| $R_{\text{free}}$<sup>g</sup> | 0.229 | 0.216 | 0.249 |
| Refined atoms |  |  |  |
| Fab | 6,852 | 3,398 | 3,341 |
| Peptide | - | 73 | 85 |
| Waters | 637 | 432 | 247 |
| $B$-values (Å<sup>2</sup>) | | | |
| Fab | 39 | 24 | 32 |
| Peptide | - | 40 | 38 |
| Waters | 42 | 32 | 30 |
| Wilson $B$-values (Å<sup>2</sup>) | 27 | 20 | 29 |
| Ramachandran statistics (%)<sup>h</sup> |  |  |  |
| Favored | 97.3 | 97.7 | 96.8 |
| Outliers | 0.5 | 0.0 | 0.4 |
| R.m.s.d. bond length (Å) | 0.007 | 0.005 | 0.007 |
| R.m.s.d. bond angles (°) | 1.12 | 0.94 | 0.95 |
| PDB ID | XXXX | XXXX | XXXX |

<sup>a</sup> Parentheses denote outer-shell statistics.
<sup>b</sup> $R_{\text{sym}} = \sum_{hkl} \sum_i |I_{hkl,i} - \langle I_{hkl} \rangle| / \sum_{hkl} \sum_i I_{hkl,i}$ and $R_{\text{pim}} = \sum_{hkl} [1/(N-1)]^{1/2} \sum_i |I_{hkl,i} - \langle I_{hkl} \rangle| / \sum_{hkl} \sum_i I_{hkl,i}$ , where $I_{hkl,i}$ is the scaled intensity of the $i$ measurement of reflection $h, k, l$ , $\langle I_{hkl} \rangle$ is the average intensity for that reflection, and $N$ is the redundancy.
<sup>c</sup> $R_{\text{pim}} = \sum_{hkl} (1/(n-1))^{1/2} \sum_i |I_{hkl,i} - \langle I_{hkl} \rangle| / \sum_{hkl} \sum_i I_{hkl,i}$ , where $n$ is the redundancy.
<sup>d</sup> $CC_{1/2}$ = Pearson Correlation Coefficient between two random half datasets.
<sup>e</sup> No. molecules for complexes refers to number of Fab molecules per asymmetric unit (ASU).
<sup>f</sup> $R_{\text{cryst}} = \sum_{hkl} |F_o - F_c| / \sum_{hkl} |F_o|$ , where $F_o$ and $F_c$ are the observed and calculated structure factors, respectively.
<sup>g</sup> $R_{\text{free}}$ was calculated as for $R_{\text{cryst}}$ , but with 5% of data excluded before refinement.
<sup>h</sup> Values are percentage of residues in the favored and outlier regions as analyzed by MolProbity (Chen et al, 2010).

**Table S3.**
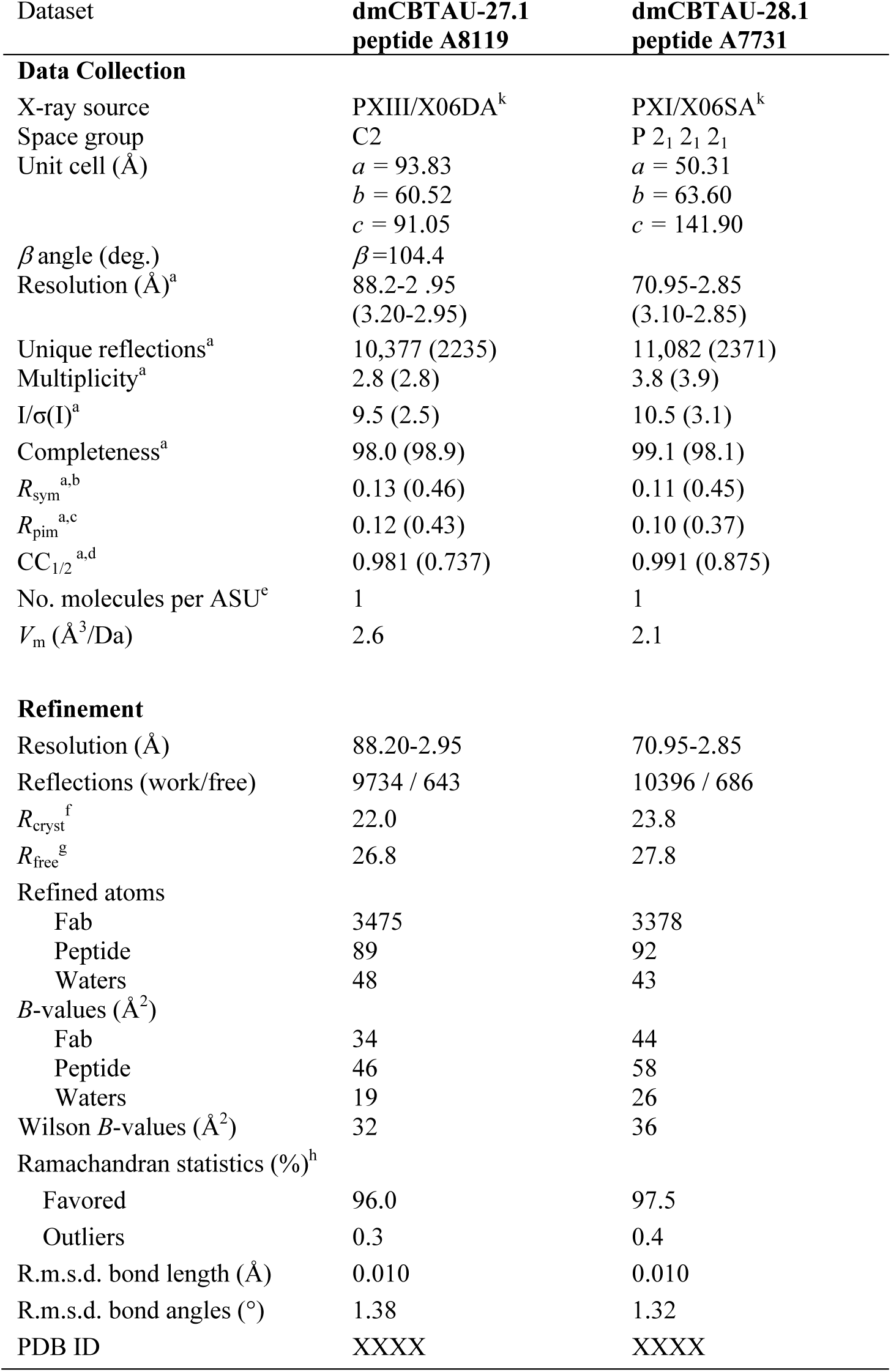
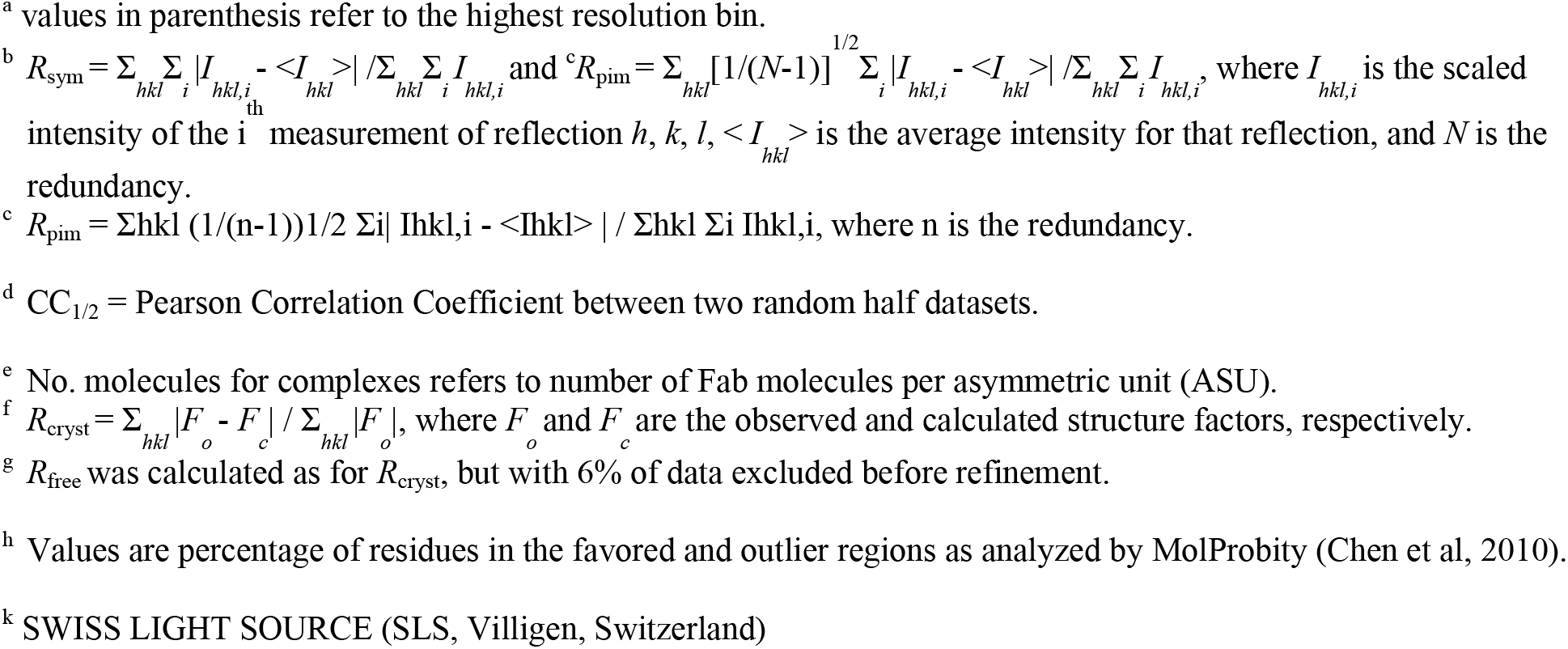
Data collection and refinement statistics for dmCBTAU-27.1 - A8119 and dmCBTAU-28.1 - A7731 complexes.

| Dataset | dmCBTAU-27.1<br>peptide A8119 | dmCBTAU-28.1<br>peptide A7731 |
| --- | --- | --- |
| **Data Collection** |  |  |
| X-ray source | PXIII/X06DA<sup>k</sup> | PXI/X06SA<sup>k</sup> |
| Space group | C2 | P 2<sub>1</sub> 2<sub>1</sub> 2<sub>1</sub> |
| Unit cell (Å) | *a* = 93.83<br>*b* = 60.52<br>*c* = 91.05 | *a* = 50.31<br>*b* = 63.60<br>*c* = 141.90 |
| $\beta$ angle (deg.) | $\beta$=104.4 | |
| Resolution (Å)<sup>a</sup> | 88.2-2.95<br>(3.20-2.95) | 70.95-2.85<br>(3.10-2.85) |
| Unique reflections<sup>a</sup> | 10,377 (2235) | 11,082 (2371) |
| Multiplicity<sup>a</sup> | 2.8 (2.8) | 3.8 (3.9) |
| I/$\sigma$(I)<sup>a</sup> | 9.5 (2.5) | 10.5 (3.1) |
| Completeness<sup>a</sup> | 98.0 (98.9) | 99.1 (98.1) |
| $R_{\text{sym}}$<sup>a,b</sup> | 0.13 (0.46) | 0.11 (0.45) |
| $R_{\text{pim}}$<sup>a,c</sup> | 0.12 (0.43) | 0.10 (0.37) |
| $CC_{1/2}$<sup>a,d</sup> | 0.981 (0.737) | 0.991 (0.875) |
| No. molecules per ASU<sup>e</sup> | 1 | 1 |
| $V_m$ (Å<sup>3</sup>/Da) | 2.6 | 2.1 |
| **Refinement** |  |  |
| Resolution (Å) | 88.20-2.95 | 70.95-2.85 |
| Reflections (work/free) | 9734 / 643 | 10396 / 686 |
| $R_{\text{cryst}}$<sup>f</sup> | 22.0 | 23.8 |
| $R_{\text{free}}$<sup>g</sup> | 26.8 | 27.8 |
| Refined atoms |  |  |
| Fab | 3475 | 3378 |
| Peptide | 89 | 92 |
| Waters | 48 | 43 |
| *B*-values (Å<sup>2</sup>) |  |  |
| Fab | 34 | 44 |
| Peptide | 46 | 58 |
| Waters | 19 | 26 |
| Wilson *B*-values (Å<sup>2</sup>) | 32 | 36 |
| Ramachandran statistics (%)<sup>h</sup> |  |  |
| Favored | 96.0 | 97.5 |
| Outliers | 0.3 | 0.4 |
| R.m.s.d. bond length (Å) | 0.010 | 0.010 |
| R.m.s.d. bond angles (°) | 1.38 | 1.32 |
| PDB ID | XXXX | XXXX |

<sup>a</sup> values in parenthesis refer to the highest resolution bin.
<sup>b</sup> $R_{\text{sym}} = \sum_{hkl} \sum_i |I_{hkl,i} - \langle I_{hkl} \rangle| / \sum_{hkl} \sum_i I_{hkl,i}$ and $R_{\text{pim}} = \sum_{hkl} [1/(N-1)]^{1/2} \sum_i |I_{hkl,i} - \langle I_{hkl} \rangle| / \sum_{hkl} \sum_i I_{hkl,i}$ , where $I_{hkl,i}$ is the scaled intensity of the $i$ measurement of reflection $h, k, l$ , $\langle I_{hkl} \rangle$ is the average intensity for that reflection, and $N$ is the redundancy.
<sup>c</sup> $R_{\text{pim}} = \sum_{hkl} (1/(n-1))^{1/2} \sum_i |I_{hkl,i} - \langle I_{hkl} \rangle| / \sum_{hkl} \sum_i I_{hkl,i}$ , where $n$ is the redundancy.
<sup>d</sup> $CC_{1/2}$ = Pearson Correlation Coefficient between two random half datasets.
<sup>e</sup> No. molecules for complexes refers to number of Fab molecules per asymmetric unit (ASU).
<sup>f</sup> $R_{\text{cryst}} = \sum_{hkl} |F_o - F_c| / \sum_{hkl} |F_o|$ , where $F_o$ and $F_c$ are the observed and calculated structure factors, respectively.
<sup>g</sup> $R_{\text{free}}$ was calculated as for $R_{\text{cryst}}$ , but with 6% of data excluded before refinement.
<sup>h</sup> Values are percentage of residues in the favored and outlier regions as analyzed by MolProbity (Chen et al, 2010).
<sup>k</sup> SWISS LIGHT SOURCE (SLS, Villigen, Switzerland)

## EXTENDED VIEW FIGURE LEGENDS

**Figure S1.**
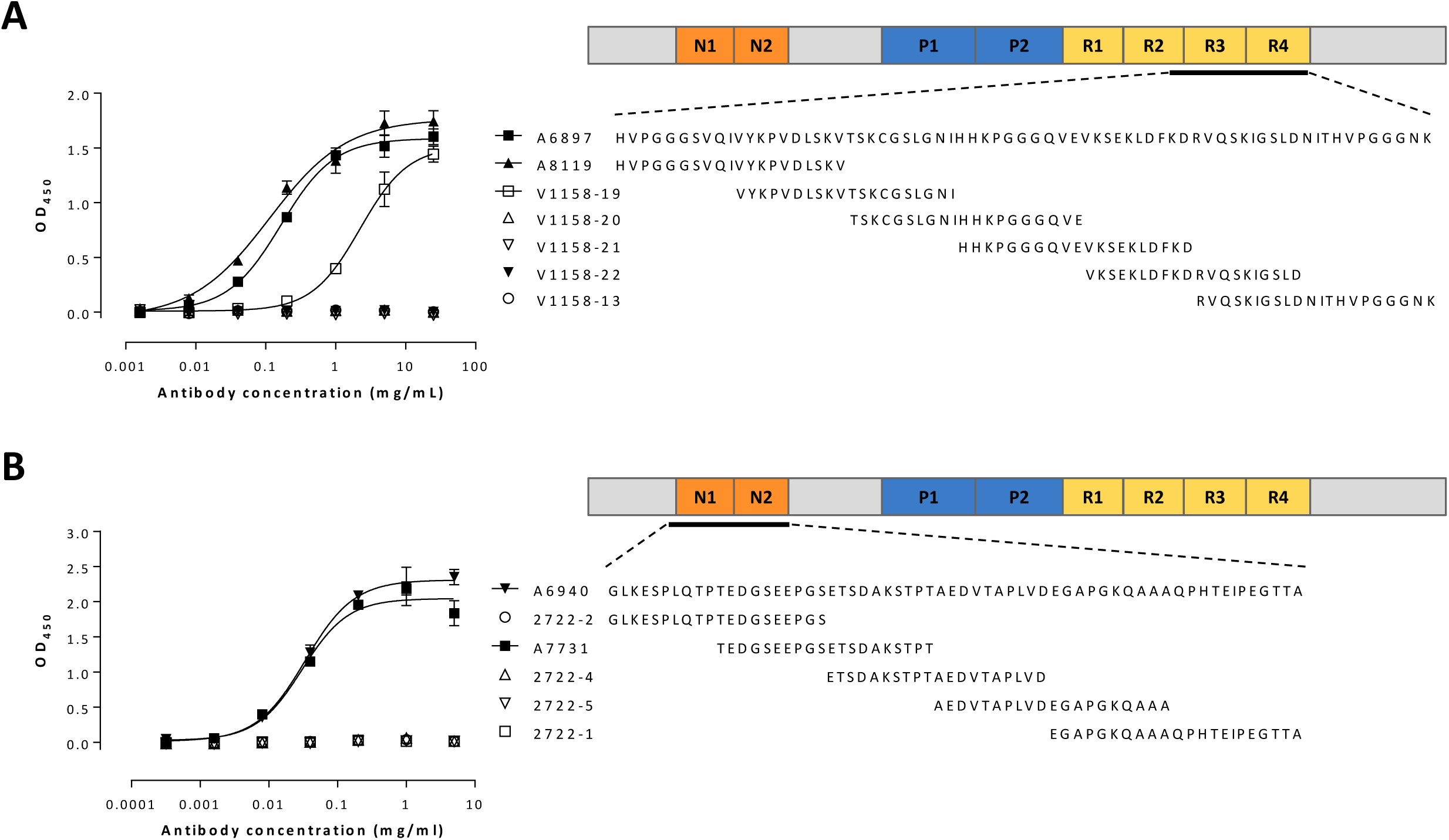
Peptide epitope mapping. Binding of CBTAU-27.1 (**A**) and CBTAU-28.1 (**B**) to tau peptides encompassing rtau residues 299-369 and 42-103, respectively, as measured by ELISA.

**Figure S2.**
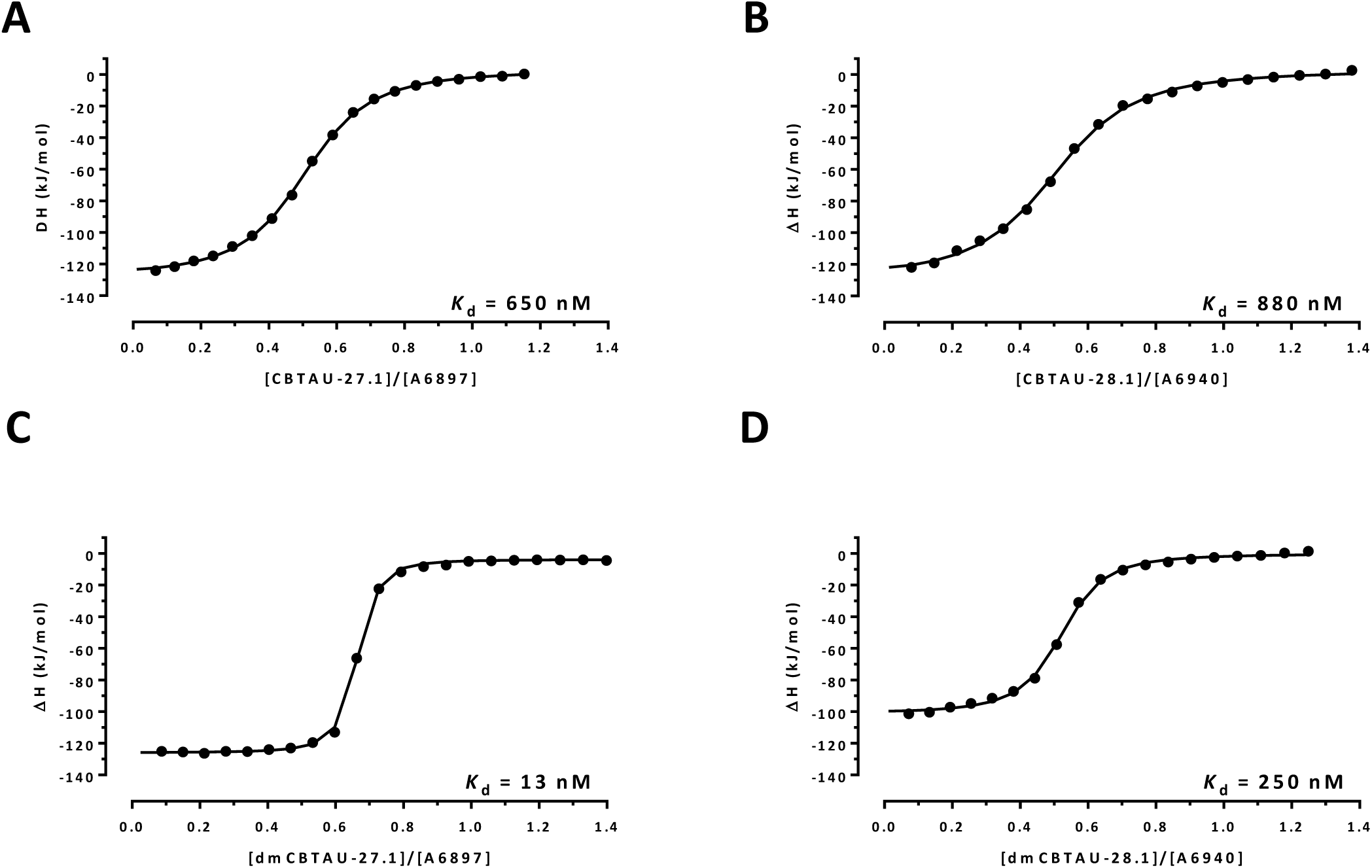
Affinity of CBTAU-27.1, CBTAU-28.1 and their affinity-matured mutants for their cognate tau peptides. Isothermal Titration Calorimetry (ITC) measurements for CBTAU-27.1 (**A**), CBTAU-28.1 (**B**), dmCBTAU-27.1 (**C**) and dmCBTAU-28.1 (**D**). Variation in enthalpy is observed following incremental addition of mAb stock to tau peptide. Continuous lines represent the best fit of experimental data assuming a single set of binding sites. Experiments were performed in PBS. Equilibrium dissociation constants (*K*_d_) are shown on the individual graphs.

**Figure S3.**
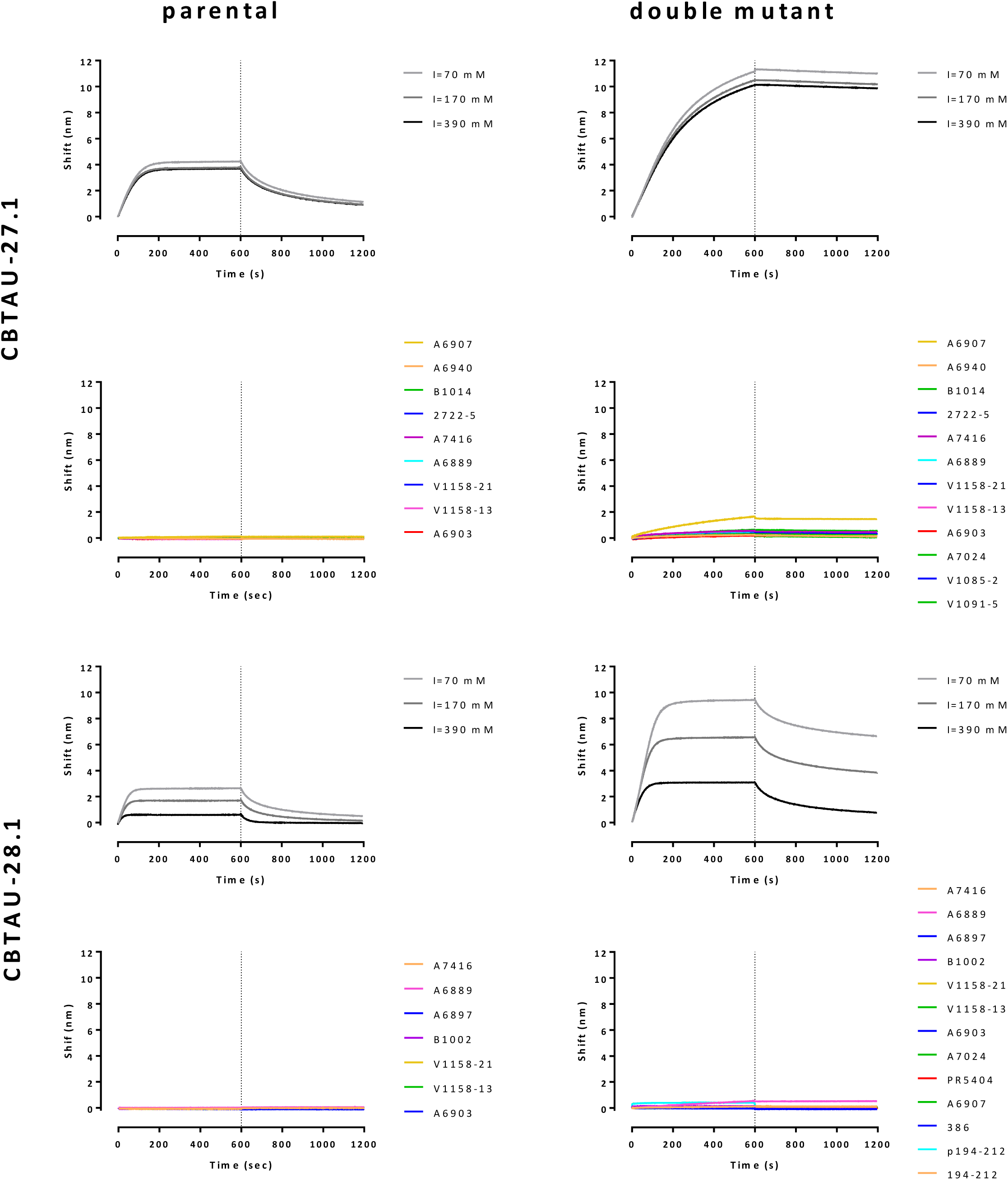
Affinity-improved antibodies dmCBTAU-27.1 and dmCBTAU-28.1 retain both the nature and specificity of the interactions of the parental antibodies with tau. Association and dissociation kinetics for the binding of CBTAU-27.1 and dmCBTAU-27.1 to peptide A6897, and of CBTAU-28.1 and dmCBTAU-28.1 to peptide A6940, were determined at different ionic strengths. In accord with the hydrophobic nature of the interaction of CBTAU-27.1 with tau, the ionic strength of the buffer did not significantly affect binding of parental antibody CBTAU-27.1 and its affinity-improved mutant. Given the charged nature of the interaction of CBTAU-28.1 with tau, binding of this antibody was significantly affected by ionic strength of the buffer and also observed for its affinity-improved mutant. For specificity, association and dissociation kinetics for the interactions of the antibodies with a panel of different tau peptides were determined. Lack of significant binding of the affinity-improved mutants to tau peptides to which the parental antibodies did not bind, and to additional peptides not containing the epitopes of the parental antibodies, indicate that the specificity was retained.

**Figure S4.**
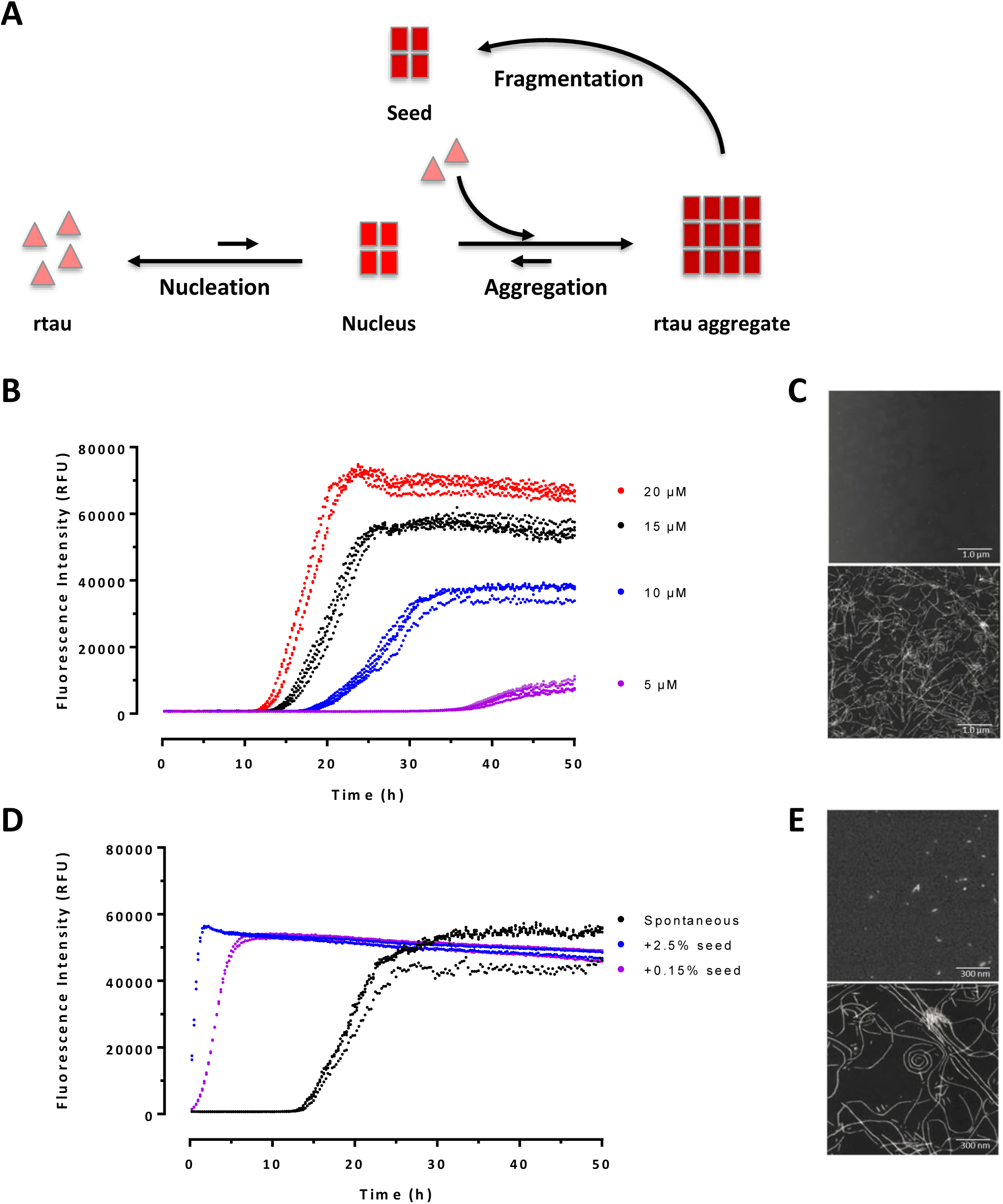
Set-up of an *in vitro* rtau aggregation assay. (**A**) Schematic representation of the nucleation-dependent polymerization process of tau aggregation. The process can be described by an initial energetically unfavorable nucleation phase followed by a fast, energetically downhill growth phase. Addition of pre-formed aggregates (“seeds”) leads to a bypass of the nucleation phase. (**B**) Kinetics for spontaneous rtau aggregation. Aggregation of rtau is induced by addition of heparin in a rTau: heparin ratio of 1:0.5 and continuously monitored by ThT fluorescence (50 μM ThT, excitation at 450 nm and emission at 482 nm) in reaction buffer. The misfolding and aggregation of rtau is a concentration-dependent process. The four independent replicates for each concentration show the high reproducibility of the assay. The obtained aggregates have PHF-like structures as assessed by (**C**) atomic force microscopy (AFM) images of a 15 µM rtau sample taken at 0 (top) and 50 (bottom) hours after initiating of aggregation. (**D**) Kinetics for spontaneous and seeded rtau aggregation in the *in vitro* aggregation assay. For seeding, 2.5 or 0.15 % (v/v) sonicated preformed aggregates were added to fresh rtau monomer solution. Addition of small amounts of pre-formed sonicated fibrillar structures leads to a bypass of the lag phase resulting in long *de novo* generated PHF-like fibrillar structures as observed by AFM. (**E**) Atomic force microscopy (AFM) images of a 2.5% seeded rtau aggregation sample taken at 0 (top) and 10 (bottom) hours after initiation of aggregation.

**Figure S5.**
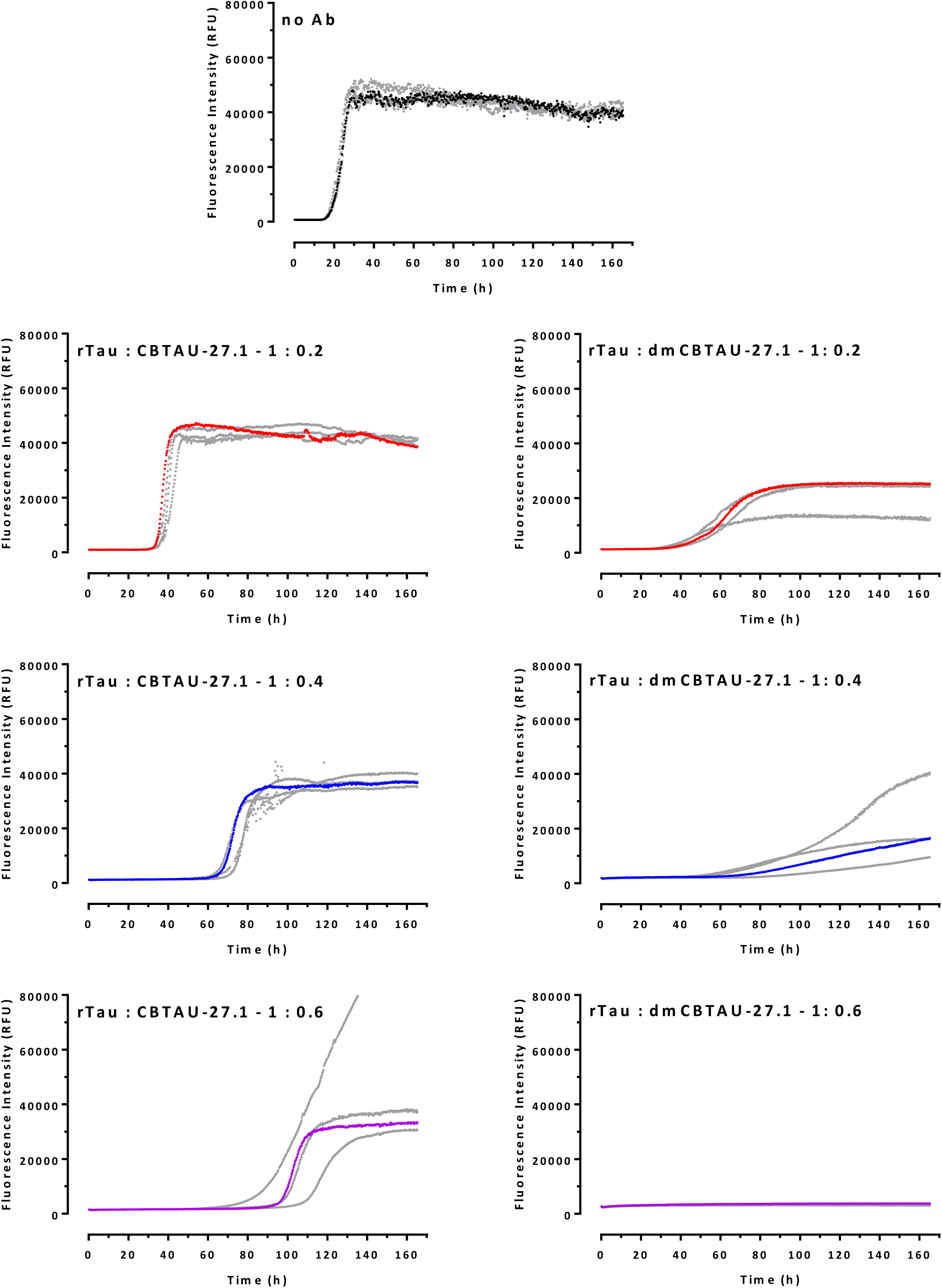
Complete dataset for *in vitro* tau aggregation in the absence or presence of CBTAU-27.1 and dmCBTAU-27.1. All four replicates for the spontaneous rtau conversion and each rtau: IgG ratio are shown in separate graphs and representative curves shown in Fig. 4A and B are indicated here in corresponding color.

**Figure S6.**
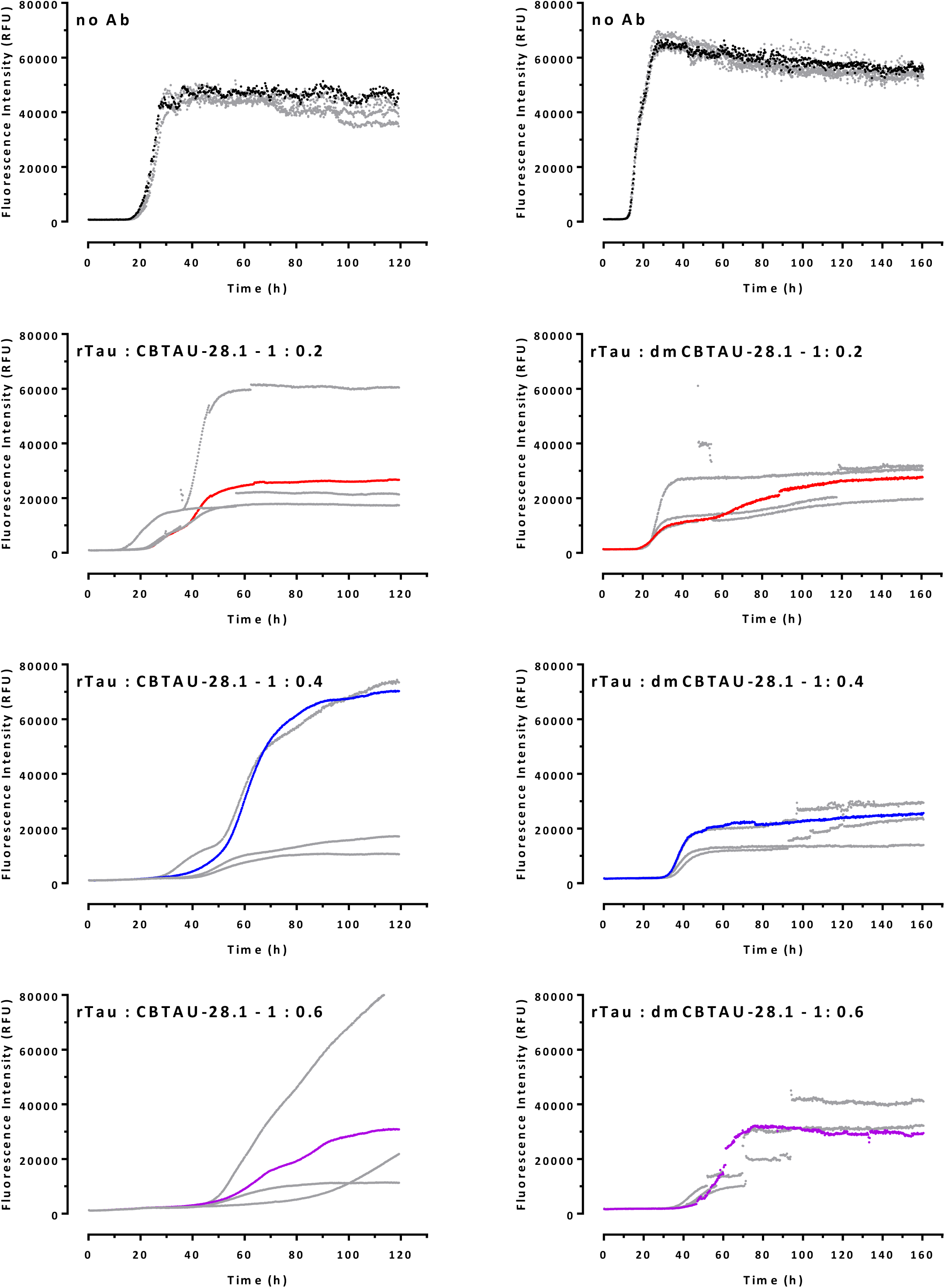
Complete dataset for *in vitro* tau aggregation in the absence or presence of CBTAU-28.1 and dmCBTAU-28.1. All four replicates for the spontaneous rtau conversion and each rtau: IgG ratio are shown in separate graphs and representative curves shown in Fig. 4E and F are indicated here in corresponding colors.

**Figure S7.**
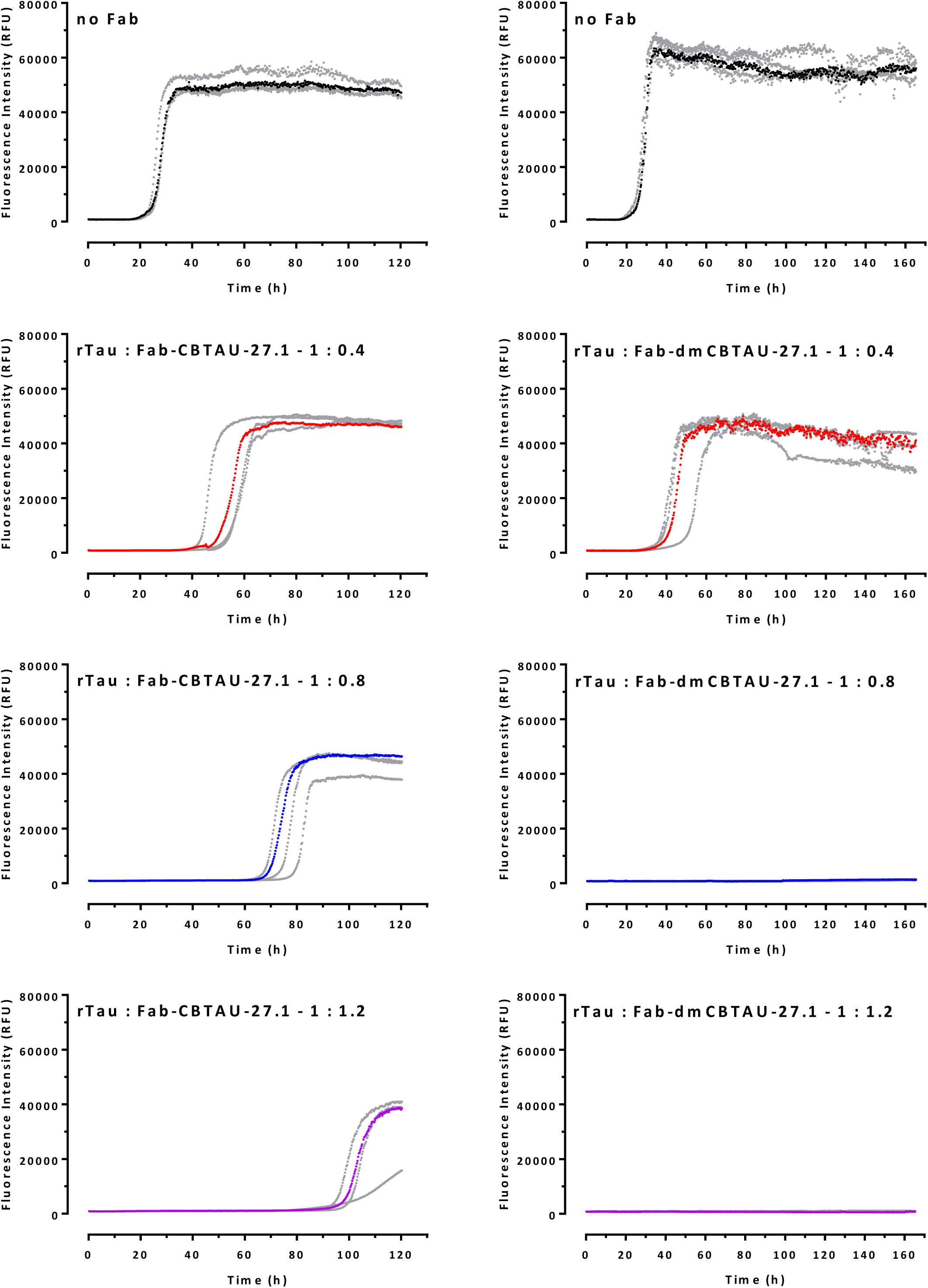
Complete dataset for *in vitro* tau aggregation in the absence or presence of Fab-CBTAU-27.1 and Fab-dmCBTAU-27.1. All four replicates for the spontaneous rtau conversion and each rtau: Fab ratio are shown in separate graphs and representative curves shown in Fig. 4C and D are indicated here in corresponding colors.

**Figure S8.**
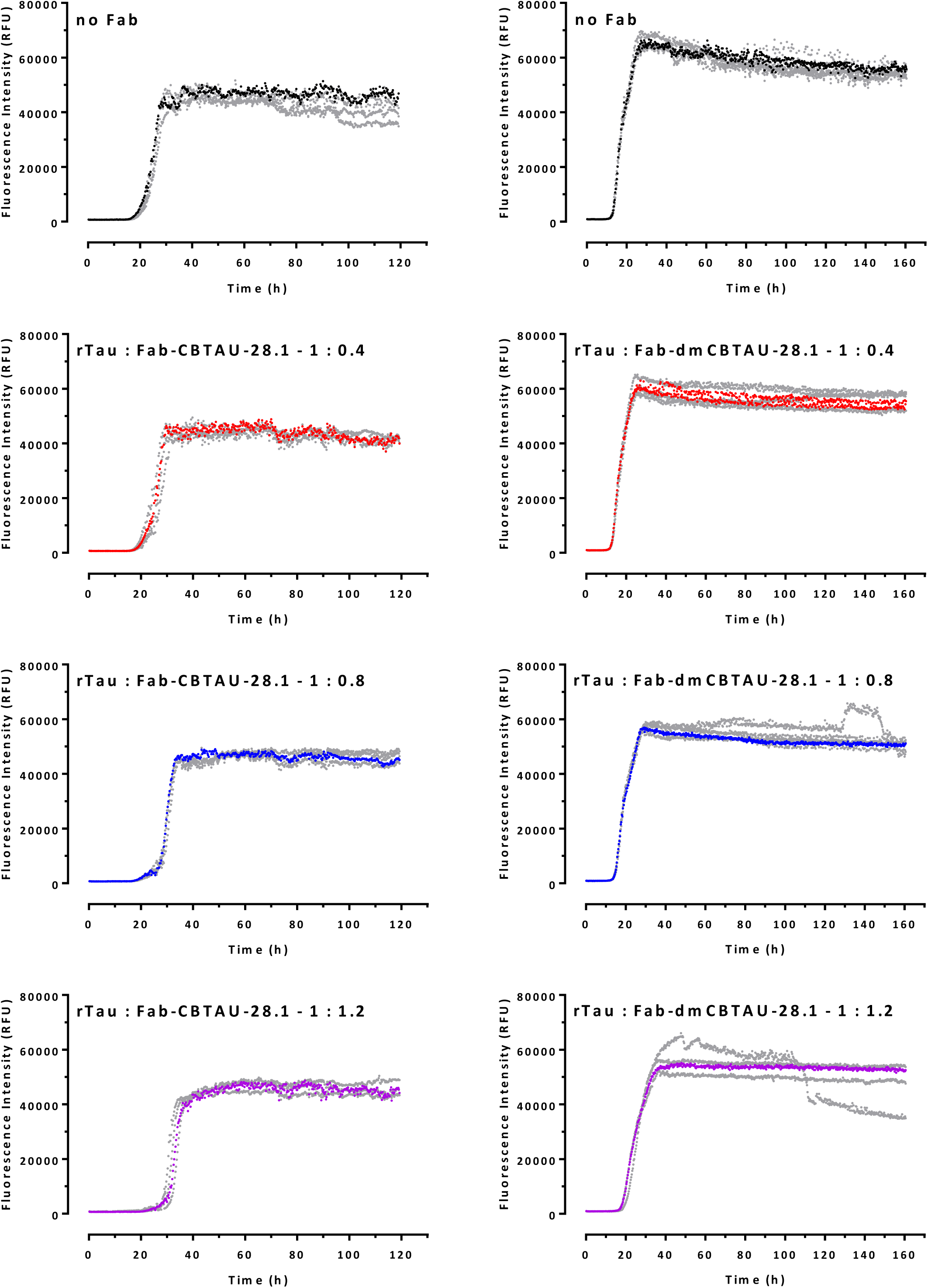
Complete dataset for *in vitro* tau aggregation in the absence or presence of Fab-CBTAU-28.1 and Fab-dmCBTAU-28.1. All four replicates for the spontaneous rtau conversion and each rtau: Fab ratio are shown in separate graphs and representative curves shown in Fig. 4F and H are indicated here in corresponding colors.

**Figure S9.**
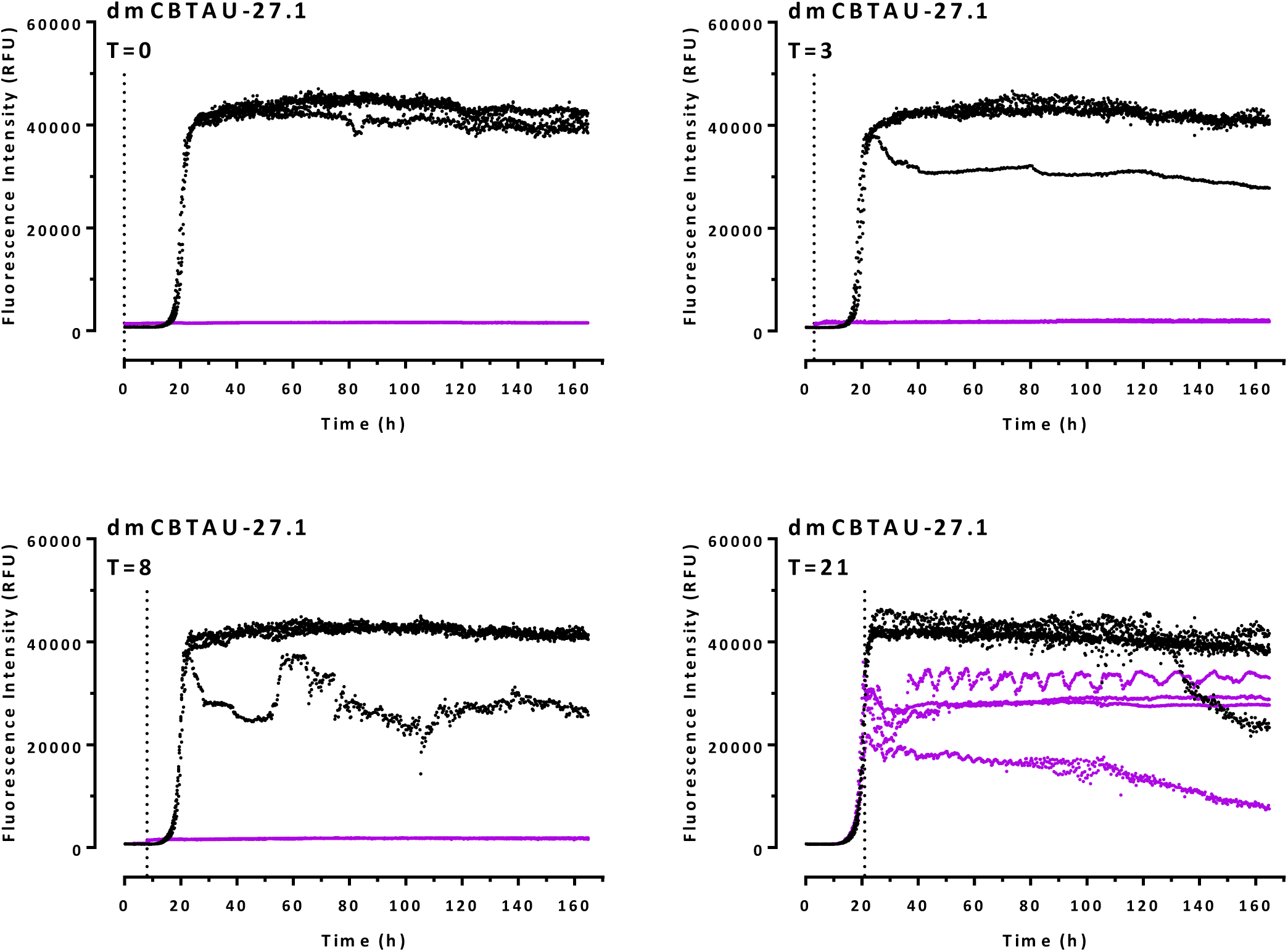
*In vitro* tau aggregation in the presence of dmCBTAU-27.1 added at different time points. *In vitro* tau aggregation (four replicates for each condition) in the presence of dmCBTAU-27.1 at a rtau:IgG ratio of 1:0.6 (purple) added at different time points (hours). For each time point, kinetics was also followed upon addition of equivalent volumes of reaction buffer (black) as control. Time points at which the antibody or reaction buffer were added are indicated by the dotted vertical line in each panel. Data indicate that CBTAU-27.1 interferes with tau aggregation by sequestering monomeric tau, also after initiation of aggregation.

**Figure S10.**
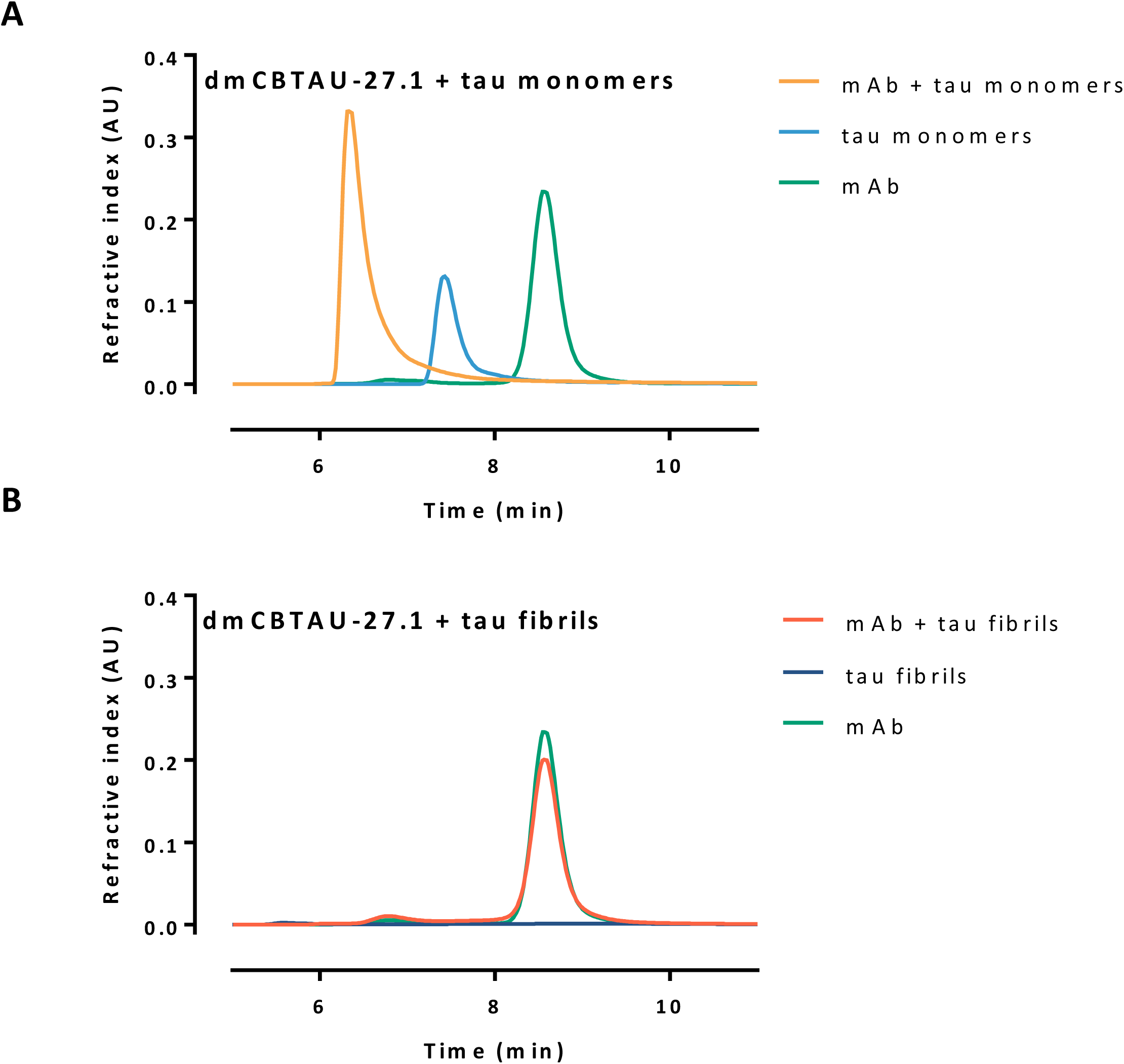
Assessment of CBTAU-27.1 binding to rtau and PHFs by SEC-MALS. SEC-MALS analysis of mixtures of antibody and rtau monomers (**A**) or fibrils (**B**), mixed in molar ratio of 1:0.6, incubated at room temperature for 15 min and subsequently centrifuged for 15 min at 20 000 g. The supernatant was taken and injected on SEC-MALS. While binding is observed between the antibody and the tau monomer, as indicated by the shift in retention time of the main peak (**A**), no tau was detected in the supernatant of the tau fibrils sample (**B**), which indicates that it sediment during centrifugation. The majority of antibody was found back in the supernatant of the mixture mAb + fibrils, indicating that the antibody did not bind to the aggregated tau.

**Figure S11.**
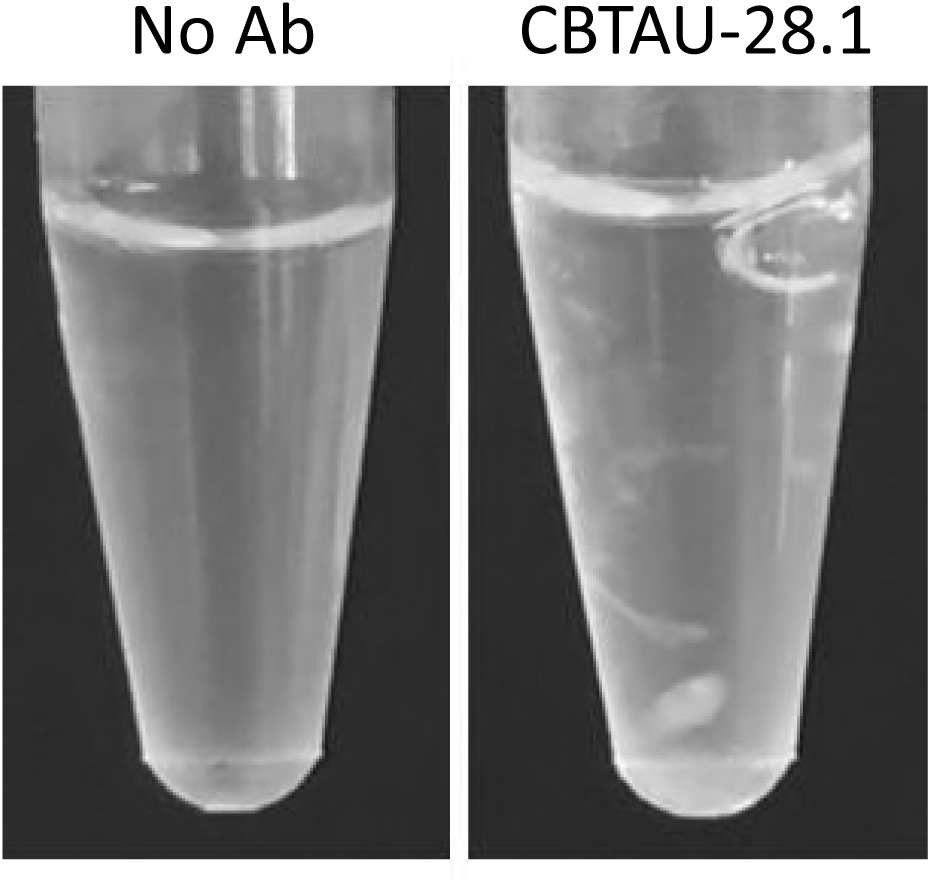
Macroscopic image of rtau fibrils generated in the absence and presence of CBTAU-28.1. Clumping indicates the capacity of CBTAU-28.1 to inducing large polymeric structures by crosslinking tau aggregates.

**Figure S12.**
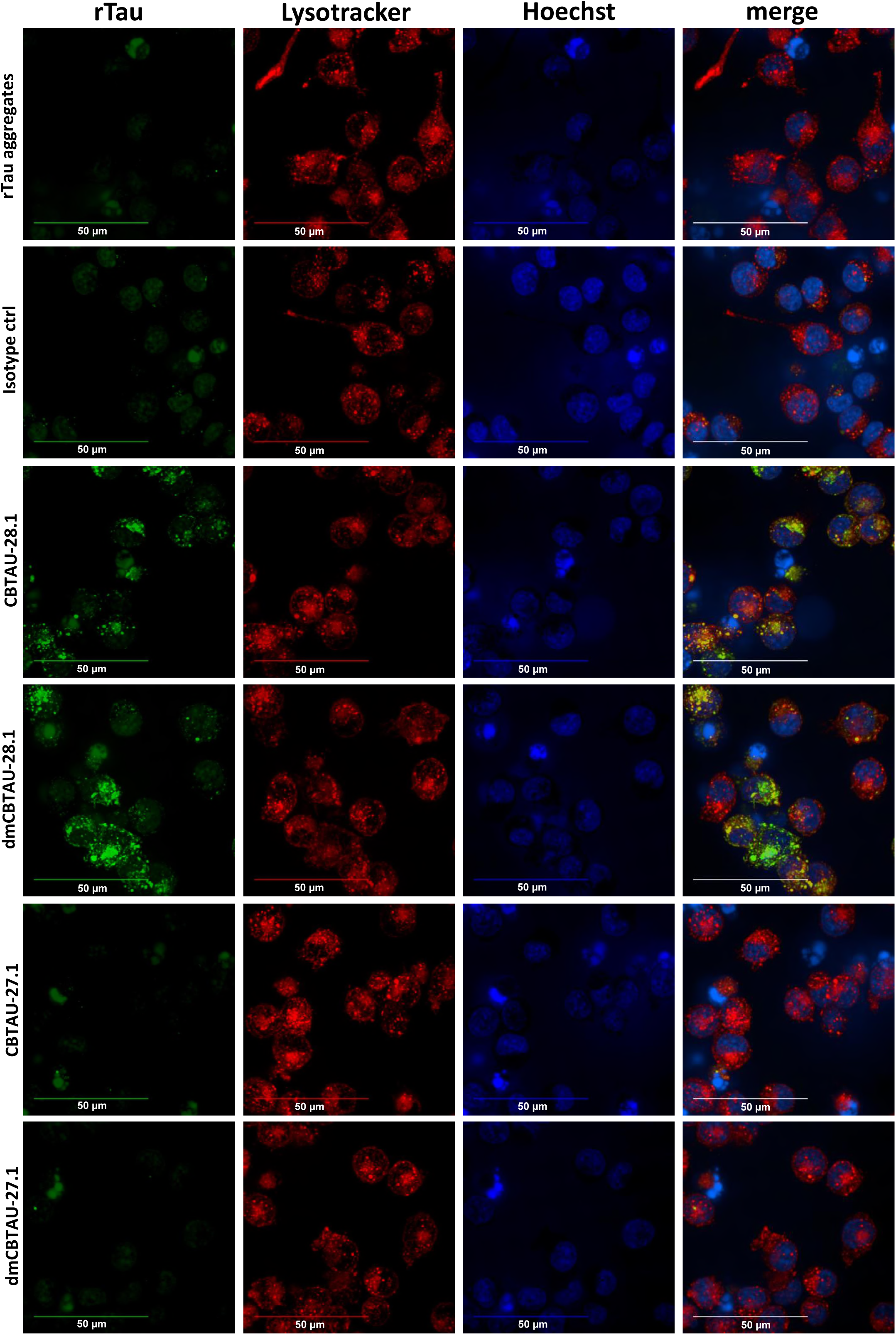
Tau aggregates are internalized by BV-2 cells and localize in cellular acidic organelles. Preformed pHrodo-Green labeled aggregated rtau/antibody immunocomplexes [chimeric CBTAU-28.1, dmCBTAU-28.1, CBTAU-27.1, dmCBTAU-27.1 and Fab fragments of these; mouse IgG1 isotype control] were incubated with BV-2 cells for 2 hours in the presence of Heparin to block antibody-independent uptake. After incubation, nuclei were stained with Hoechst (blue) and the acidic cellular compartment with LysoTracker Red dye. Live-cell imaging revealed intracellular puncta of pHrodo-Green labeled rtau aggregates inside the cells that were incubated with CBTAU-28.1 and dmCBTAU-28.1, but not with CBTAU-27.1 and dmCBTAU-27.1 or isotype control. Moreover, intracellular rtau often colocalized with LysoTracker Red thus suggesting presence of rtau in the acidic cellular compartment. Images represent maximum intensity projections of a 20 planes Z-stack (0.5 μm planes) acquired with a 63x water immersion objective.

## REFERENCES

Afonine PV, Grosse-Kunstleve RW, Echols N, Headd JJ, Moriarty NW, Mustyakimov M, Terwilliger TC, Urzhumtsev A, Zwart PH, Adams PD (2012) Towards automated crystallographic structure refinement with phenix.refine. *Acta crystallographica Section D*, Biological crystallography 68: 352–367

Apetri AC, Vanik DL, Surewicz WK (2005) Polymorphism at residue 129 modulates the conformational conversion of the D178N variant of human prion protein 90-231. Biochemistry 44: 15880–15888

Arevalo JH, Stura EA, Taussig MJ, Wilson IA (1993) Three-dimensional structure of an anti-steroid Fab’ and progesterone-Fab’ complex. J Mol Biol 231: 103–118

Billingsley ML, Kincaid RL (1997) Regulated phosphorylation and dephosphorylation of tau protein: effects on microtubule interaction, intracellular trafficking and neurodegeneration. Biochem J 323 ( **Pt 3**): 577–591

Braak H, Braak E (1991) Neuropathological stageing of Alzheimer-related changes. Acta Neuropathol 82: 239–259

Bright J, Hussain S, Dang V, Wright S, Cooper B, Byun T, Ramos C, Singh A, Parry G, Stagliano N, Griswold-Prenner I (2015) Human secreted tau increases amyloid-beta production. Neurobiol Aging 36: 693–709

Chen VB, Arendall WB, 3rd, Headd JJ, Keedy DA, Immormino RM, Kapral GJ, Murray LW, Richardson JS, Richardson DC (2010) MolProbity: all-atom structure validation for macromolecular crystallography. Acta crystallographica Section D, Biological crystallography 66: 12–21

Chen X, Zhao, Y., Harlos, K., Snir, O., Sollid, L.M. (2017) Crystal structure of anti-gliadin 1002-1E03 Fab fragment in complex of peptide PLQPEQPFP. DOI 102210/pdb5ijk/pdb

Cleveland DW, Hwo SY, Kirschner MW (1977a) Physical and chemical properties of purified tau factor and the role of tau in microtubule assembly. J Mol Biol 116: 227–247

Cleveland DW, Hwo SY, Kirschner MW (1977b) Purification of tau, a microtubule-associated protein that induces assembly of microtubules from purified tubulin. J Mol Biol 116: 207–225

Concepcion J, Witte K, Wartchow C, Choo S, Yao D, Persson H, Wei J, Li P, Heidecker B, Ma W, Varma R, Zhao LS, Perillat D, Carricato G, Recknor M, Du K, Ho H, Ellis T, Gamez J, Howes M, Phi-Wilson J, Lockard S, Zuk R, Tan H (2009) Label-free detection of biomolecular interactions using BioLayer interferometry for kinetic characterization. Comb Chem High Throughput Screen 12: 791–800

Crespo R, Rocha FA, Damas AM, Martins PM (2012) A generic crystallization-like model that describes the kinetics of amyloid fibril formation. J Biol Chem 287: 30585–30594

Daebel V, Chinnathambi S, Biernat J, Schwalbe M, Habenstein B, Loquet A, Akoury E, Tepper K, Muller H, Baldus M, Griesinger C, Zweckstetter M, Mandelkow E, Vijayan V, Lange A (2012) beta-Sheet core of tau paired helical filaments revealed by solid-state NMR. J Am Chem Soc 134: 13982–13989

Dehmelt L, Halpain S (2005) The MAP2/Tau family of microtubule-associated proteins. Genome Biol 6: 204

Emsley P, Lohkamp B, Scott WG, Cowtan K (2010) Features and development of Coot. Acta Crystallogr D Biol Crystallogr 66: 486–501

Fitzpatrick AWP, Falcon B, He S, Murzin AG, Murshudov G, Garringer HJ, Crowther RA, Ghetti B, Goedert M, Scheres SHW (2017) Cryo-EM structures of tau filaments from Alzheimer’s disease. Nature 547: 185–190

Frost B, Jacks RL, Diamond MI (2009) Propagation of tau misfolding from the outside to the inside of a cell. J Biol Chem 284: 12845–12852

Ginhoux F, Lim S, Hoeffel G, Low D, Huber T (2013) Origin and differentiation of microglia. Front Cell Neurosci 7: 45

Goedert M, Spillantini MG, Jakes R, Rutherford D, Crowther RA (1989) Multiple isoforms of human microtubule-associated protein tau: sequences and localization in neurofibrillary tangles of Alzheimer’s disease. Neuron 3: 519–526

Gorny MK, Sampson J, Li H, Jiang X, Totrov M, Wang XH, Williams C, O’Neal T, Volsky B, Li L, Cardozo T, Nyambi P, Zolla-Pazner S, Kong XP (2011) Human anti-V3 HIV-1 monoclonal antibodies encoded by the VH5-51/VL lambda genes define a conserved antigenic structure. PloS one 6: e27780

Guo JL, Lee VM (2011) Seeding of normal Tau by pathological Tau conformers drives pathogenesis of Alzheimer-like tangles. J Biol Chem 286: 15317–15331

Guo JL, Narasimhan S, Changolkar L, He Z, Stieber A, Zhang B, Gathagan RJ, Iba M, McBride JD, Trojanowski JQ, Lee VM (2016) Unique pathological tau conformers from Alzheimer’s brains transmit tau pathology in nontransgenic mice. J Exp Med 213: 2635–2654

Harper JD, Lansbury PT, Jr. (1997) Models of amyloid seeding in Alzheimer’s disease and scrapie: mechanistic truths and physiological consequences of the time-dependent solubility of amyloid proteins. Annu Rev Biochem 66: 385–407

Holmes BB, Furman JL, Mahan TE, Yamasaki TR, Mirbaha H, Eades WC, Belaygorod L, Cairns NJ, Holtzman DM, Diamond MI (2014) Proteopathic tau seeding predicts tauopathy in vivo. Proc Natl Acad Sci USA 111: E4376–4385

Kabsch W (2010) Xds. Acta Crystallogr Sect D Biol Crystallogr 66: 125–132

Kfoury N, Holmes BB, Jiang H, Holtzman DM, Diamond MI (2012) Trans-cellular propagation of Tau aggregation by fibrillar species. J Biol Chem 287: 19440–19451

Kontsekova E, Zilka N, Kovacech B, Skrabana R, Novak M (2014) Identification of structural determinants on tau protein essential for its pathological function: novel therapeutic target for tau immunotherapy in Alzheimer’s disease. Alzheimer’s Res & Ther 6: 45

Kramer RA, Cox F, van der Horst M, van der Oudenrijn S, Res PC, Bia J, Logtenberg T, de Kruif J (2003) A novel helper phage that improves phage display selection efficiency by preventing the amplification of phages without recombinant protein. Nucleic Acids Res 31: e59

Laskowski RA, MacArthur, M.W., Moss, D. S. & Thornton, J.M (1993) PROCHECK: a program to check the stereochemicai quality of protein structures. J Appl Cryst 26: 283–291

Lee VM, Balin BJ, Otvos L, Jr., Trojanowski JQ (1991) A68: a major subunit of paired helical filaments and derivatized forms of normal Tau. Science 251: 675–678

Lee VM, Goedert M, Trojanowski JQ (2001) Neurodegenerative tauopathies. Annu Rev Neurosci 24: 1121–1159

Li W, Lee VM (2006) Characterization of two VQIXXK motifs for tau fibrillization in vitro. Biochemistry 45: 15692–15701

Liao HX, Bonsignori M, Alam SM, McLellan JS, Tomaras GD, Moody MA, Kozink DM, Hwang KK, Chen X, Tsao CY, Liu P, Lu X, Parks RJ, Montefiori DC, Ferrari G, Pollara J, Rao M, Peachman KK, Santra S, Letvin NL, Karasavvas N, Yang ZY, Dai K, Pancera M, Gorman J, Wiehe K, Nicely NI, Rerks-Ngarm S, Nitayaphan S, Kaewkungwal J, Pitisuttithum P, Tartaglia J, Sinangil F, Kim JH, Michael NL, Kepler TB, Kwong PD, Mascola JR, Nabel GJ, Pinter A, Zolla-Pazner S, Haynes BF (2013) Vaccine induction of antibodies against a structurally heterogeneous site of immune pressure within HIV-1 envelope protein variable regions 1 and 2. Immunity 38: 176–186

Mandelkow E, von Bergen M, Biernat J, Mandelkow EM (2007) Structural principles of tau and the paired helical filaments of Alzheimer’s disease. Brain pathology 17: 83–90

McCoy AJ, Grosse-Kunstleve RW, Storoni LC, Read RJ (2005) Likelihood-enhanced fast translation functions. Acta Crystallogr Sect D Biol Crystallogr 61: 458–464

Ofek G, Zirkle B, Yang Y, Zhu Z, McKee K, Zhang B, Chuang GY, Georgiev IS, O’Dell S, Doria-Rose N, Mascola JR, Dimitrov DS, Kwong PD (2014) Structural basis for HIV-1 neutralization by 2F5-like antibodies m66 and m66.6. J Virol 88: 2426–2441

Pascual G, Wadia JS, Zhu X, Keogh E, Kukrer B, van Ameijde J, Inganas H, Siregar B, Perdok G, Diefenbach O, Nahar T, Sprengers I, Koldijk MH, der Linden EC, Peferoen LA, Zhang H, Yu W, Li X, Wagner M, Moreno V, Kim J, Costa M, West K, Fulton Z, Chammas L, Luckashenak N, Fletcher L, Holland T, Arnold C, Anthony Williamson R, Hoozemans JJ, Apetri A, Bard F, Wilson IA, Koudstaal W, Goudsmit J (2017) Immunological memory to hyperphosphorylated tau in asymptomatic individuals. Acta Neuropathol 133: 767–783

Sanchez C, Diaz-Nido J, Avila J (2000) Phosphorylation of microtubule-associated protein 2 (MAP2) and its relevance for the regulation of the neuronal cytoskeleton function. Prog Neurobiol 61: 133–168

Surewicz WK, Jones EM, Apetri AC (2006) The emerging principles of mammalian prion propagation and transmissibility barriers: Insight from studies in vitro. Acc Chem Res 39: 654–662

Thal DR, Rub U, Orantes M, Braak H (2002) Phases of A beta-deposition in the human brain and its relevance for the development of AD. Neurology 58: 1791–1800

Uchihara T (2007) Silver diagnosis in neuropathology: principles, practice and revised interpretation. Acta Neuropathol 113: 483–499

Umeda T, Eguchi H, Kunori Y, Matsumoto Y, Taniguchi T, Mori H, Tomiyama T (2015) Passive immunotherapy of tauopathy targeting pSer413-tau: a pilot study in mice. Ann Clin Transl Neurol 2: 241–255

Vagin A,. Teplyakov, A. (1997) MOLREP: an Automated Program for Molecular Replacement J Appl Cryst 30: 1022–1025.

Vagin AA, Steiner RA, Lebedev AA, Potterton L, McNicholas S, Long F, Murshudov GN (2004) REFMAC5 dictionary: organization of prior chemical knowledge and guidelines for its use. Acta Crystallogr Sect D Biol Crystallogr 60: 2184–2195

von Bergen M, Friedhoff P, Biernat J, Heberle J, Mandelkow EM, Mandelkow E (2000) Assembly of tau protein into Alzheimer paired helical filaments depends on a local sequence motif ((306)VQIVYK(311)) forming beta structure. Proc Natl Acad Sci USA 97: 5129–5134

Weingarten MD, Lockwood AH, Hwo SY, Kirschner MW (1975) A protein factor essential for microtubule assembly. Proc Natl Acad Sci USA 72: 1858–1862

Wu JW, Herman M, Liu L, Simoes S, Acker CM, Figueroa H, Steinberg JI, Margittai M, Kayed R, Zurzolo C, Di Paolo G, Duff KE (2013) Small misfolded Tau species are internalized via bulk endocytosis and anterogradely and retrogradely transported in neurons. J Biol Chem 288: 1856–1870

Yanamandra K, Kfoury N, Jiang H, Mahan TE, Ma S, Maloney SE, Wozniak DF, Diamond MI, Holtzman DM (2013) Anti-tau antibodies that block tau aggregate seeding in vitro markedly decrease pathology and improve cognition in vivo. Neuron 80: 402–414

Yanamandra K, Patel TK, Jiang H, Schindler S, Ulrich JD, Boxer AL, Miller BL, Kerwin DR, Gallardo G, Stewart F, Finn MB, Cairns NJ, Verghese PB, Fogelman I, West T, Braunstein J, Robinson G, Keyser J, Roh J, Knapik SS, Hu Y, Holtzman DM (2017) Anti-tau antibody administration increases plasma tau in transgenic mice and patients with tauopathy. Sci Transl Med 9

